# A family of fitness landscapes modeled through gene regulatory networks

**DOI:** 10.1101/2021.12.03.471063

**Authors:** Chia-Hung Yang, Samuel V. Scarpino

## Abstract

Over 100 years, Fitness landscapes have been a powerful metaphor for understanding the evolution of biological systems. These landscapes describe how genotypes are connected to each other and are related according to relative fitness. Despite the high dimensionality of such real-world landscapes, empirical studies are often limited in their ability to quantify the fitness of different genotypes beyond point mutations, while theoretical works attempt statistical/mechanistic models to reason the overall landscape structure. However, most classical fitness landscape models overlook an instinctive constraint that genotypes leading to the same phenotype almost certainly share the same fitness value, since the information of genotype-phenotype mapping is rarely incorporated. Here, we investigate fitness landscape models through the lens of Gene Regulatory Networks (GRNs), where the regulatory products are computed from multiple genes and collectively treated as the phenotypes. With the assumption that regulatory mediators/products exhibit binary states, we prove topographical features of GRN fitness landscape models such as accessibility and connectivity insensitive to the choice of the fitness function. Furthermore, using graph theory, we deduce a mesoscopic structure underlying GRN fitness landscape models that retains necessary information for evolutionary dynamics with minimal complexity. We also propose an algorithm to construct such a mesoscopic backbone which is more efficient than the brute-force approach. Combined, this work provides mathematical implications for fitness landscape models with high-dimensional genotype-phenotype mapping, yielding the potential to elucidate empirical landscapes and their resulting evolutionary processes in a manner complementary to existing computational studies.

## 1 Introduction

Since its first introduction by Wright [1], the concept of fitness landscapes has grown and matured into a widely studied cornerstone in evolutionary biology [2, 3, 4]. A fitness landscape consists of a space of genotypes which are mutually accessible through mutations, and a fitness value associated with each genotype that quantifies its evolutionary “success”. In the common metaphor, evolution of a population is depicted as a trajectory wondering on the fitness landscape. The topography of a fitness landscape hence sheds light on various evolutionary perspectives, including constraints of adaptation [5, 6, 7, 8], sources in post-zygotic speciation [9, 10], (dis)advantages of sex and recombination [11, 12, 13], and the repeatability of evolutionary histories [14, 15, 16, 17].

Over the past three decades, empirical work has investigated specific fitness landscapes for various organisms, ranging from bacteria [18, 19, 20], fungi [21, 22], to fruit fly [23]. The underlying genotypic space of an empirical fitness landscape is typically constructed from a wild type genotype and a collection of mutations of interest, such that the binary occurrence of each mutation on the wild type span through a hypercube of mutant genotypes. The fitness, or a proxy of fitness, is then experimentally measured for each mutant genotype. While the number of focal mutations was fairly limited in these early studies, modern sequencing techniques and high-throughput analyses enable a more comprehensive search of genotypes on the sequence level for the HIV-1 virus [24, 25], yeast [26], *E. coli* [27], jellyfish [28], cancer [29], human stem cells [30], and in biochemical contents [31, 32, 33, 34, 35]. The thereof navigable genotypes can be conceptualized as a graph termed the genotype network [36], where two nodes (genotypes) are connected if they only differ by a single mutation.

Complementarily, several models of fitness landscapes have been proposed earlier for illuminating insights of evolutionary phenomena and later to be compared against empirical landscapes. A minimal ‘house of cards’ (HoC) model [37, 38] assumes that the fitness value of each genotype is an independent sample from a given probabilistic distribution. Kauffman and Weinberger [39] introduced the NK model that forces each locus to interact with a fixed number of other loci, where a genotype’s fitness becomes the sum of fitness contribution of every interaction group, which is again a random sample drawn from some distribution. Furthermore, the rough Mount Fuji model [40, 41] combines the HoC landscape with an additional field penalizing a genotype’s Hamming distance away from a referenced genotype with the optimal fitness. On the other extreme, Fisher’s geometric model [14, 42, 43] assumes that mutations have additive effects on a few phenotypes, and it transforms fitness into a continuous and non-linear landscape over a lower-dimensional phenotypic space.

Both empirical and modeled fitness landscapes in the literature reveal several prominent topographical features. First, fitness landscapes are thought to be “rugged” instead of a smooth surface. The degree of ruggedness can be assessed via a variety of measures, such as the roughness of slope ratio [44, 45] and the number of local fitness maxima [46], which are found strongly correlated with each others [4]. Empirical studies reported moderate ruggedness in the observed fitness landscapes [19, 18, 22, 20, 47], which is less intensive than the HoC model and comparable to a fine-tuned NK model or rough Mount Fuji model [4]. Second, fitness landscapes reveal mutational trajectories from one genotype to another where the fitness is non-dreasing, which implicates the accessibility (typically to a fitness optimum) of the landscapes. Despite that a majority of trajectories do not feature a progressive fitness, accessible trajectories have been uncovered in empirical landscapes [48, 18, 49, 50, 51]. In addition, whereas the inaccessible region in the HoC model expands when being distant from the fitness optimum [52], other models are shown to have definite accessible trajectories with a high genotypic dimensionality [53, 54].

Fitness landscape models span across the genotypic and phenotypic level, yet neither one alone is a sufficient determinant of a landscape’s topography. On the one hand, correspondence between genotypes and their fitness is rather indirect; hypothetically, two genotypes leading to the same phenotype are deemed to show the same fitness. On the other hand, mutations occur on the genotypic level, so even two genotypes with the identical phenotypic behavior are not guaranteed to have equivalent local neighborhoods in a fitness landscape. An ideal fitness landscape model shall therefore incorporate correlation between the two, i.e., the genotype-phenotype mapping. Determining the resultant phenotypes from the genotypes is nevertheless a rather complex “computational” procedure [55]: Information of inheritance is processed through various mechanisms including regulatory genetic interactions, metabolic pathways, chemical stabilization under thermodynamics and organism development to eventually manifest physiological traits. Existing fitness landscape models with genotype-phenotype mapping include relating the folded structure of RNA sequences and using its stability or affinity as a fitness proxy [56, 57, 58, 59], modeling a network of molecular/genetic pathways whose resultant expression pattern or metabolic homeostasis determines the fitness [60, 61, 62, 63, 64], and integrating modular mutational effects at different loci in Fisher’s geometric model [65]. However, the increasing complexity of genotype-phenotype mapping makes analytical predictions challenging, and the fitness landscape models commonly become computationally intensive to study.

Here, we model the genotype-phenotype mapping through the pathway framework of gene regulatory networks (GRNs), where mechanistic knowledge of how phenotypes are computed from genotypes is encoded in the GRN (see [66, 67] for a more formal introduction to the pathway framework). To study the fitness landscape induced by GRN evolution, we integrate the pathway framework into a family of fitness landscape models where the fitness value is uniquely determined by the phenotype corresponding to the regulatory outcome of a genotype. For a fitness landscape of GRNs, we first establish two of its topographical features that (a) GRNs with the same phenotype are themselves connected in the underlying genotypic space, i.e., the genotype network, and (b) there exist accessible trajectories between GRNs of any two close enough phenotypes. Second, utilizing the idea of symmetries and automorphisms in the genotype network, our thorough analysis coarse-grains GRNs into groups where GRNs in the same group have equivalent roles in the fitness landscapes. Extracting each group of GRNs by a single representative then strikingly reduces the complexity of the landscape. Third, we propose an algorithm to construct the underlying space of the fitness landscape of GRNs, which is arguably more efficient than the conventional brute-force approach since it incrementally builds up the equivalent groups of GRNs and simultaneously computes their mutational connection.

## 2 Methods

Here we introduce a family of fitness landscape models where the genotype-phenotype mapping is constructed from regulatory genetic interactions. We first summarize a modeling framework of GRNs proposed in our previous work [66, 67], termed the pathway framework, and then fitness landscape models of GRNs built upon the pathway framework are formally introduced.

### 2.1 Pathway framework of GRNs

In a nutshell, the pathway framework of GRNs [66, 67] abstracts genotypes on the level of their gene expression behavior: The allele of each locus is represented by the pair of its transcription activator and the protein product. Based on the activator/product pairs of a given set of genes, the transcriptional regulatory interactions among them can be deduced where one gene’s expression product also corresponds to the activator of another. More importantly, the pathway framework models the signals of external environmental stimuli propagating and cascading through these regulatory genetic interactions to develop the chemical state of protein activators/products, and the resultant phenotype consists of the collective state of a pre-specified set of proteins. For more details about the pathway framework of GRNs, see [66, 67] for references.

The general idea of the pathway framework can be flexibly adopted and adjusted for different modeling scenarios. In this work, we restrict the pathway framework into a few further assumptions when building the family of fitness landscape models: (a) We consider a fixed set of genes underlying the genotypes, i.e., gene duplication and deletion events are excluded in the resultant fitness landscape models. (b) We also assume a fixed underlying collection of proteins that can possibly exist in the organism, and only a subset of them relates to the developed phenotype. (c) We will consider the cases where a gene’s expression behavior can only be activated by one specific protein, and it generates a single protein product. (d) We assume that the associated chemical state of each protein is modeled as a Boolean/binary variable (present or absent), and external environmental signals stimulate the existence of specific proteins in the organism.

### 2.2 Fitness landscape of GRNs under the pathway framework

Let Γ and Ω be the fixed, underlying collection of loci and proteins respectively. A genotype is represented by its GRN *g* such that every locus *γ* ∈Γ is associated with a pair of protein activator/product *e*_*g*_(*γ*) = (*u, v*), *u, v* ∈ Ω. Equivalently, any GRN is a directed graph with |Ω| nodes labeled by the proteins Ω and |Γ| edges labeled by the loci Γ. In the rest of this paper, we will use the terminology “source/target node of edge *γ*” to interexchangeably refer to the protein activator/product of locus *γ*. We also write 𝒢 to be the set of all GRNs with the underlying loci Γ and proteins Ω.

The backbone of a fitness landscape of GRNs, i.e., the genotype network, is an undirected network of networks consisting of mutational relationship between the GRNs. Let *G* be the genotype network, and we denote its mega-nodes by *V*(*G*) = 𝒢 and its edges by *E*(*G*). There is an edge (*g*_1_, *g*_2_) ∈ *E*(*G*) between any GRNs *g*_1_, *g*_2_ ∈ 𝒢 when they only differ by the allele of a single locus 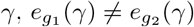. In other words, *g*_1_ and *g*_2_ are connected in *G* when they can be transformed into each other through one edge rewiring.

Furthermore, we write *x*_*ω*_ to be the binary state of protein *ω* ∈ Ω, where *x*_*ω*_ = 1 indicates the presence of *ω* and *x*_*ω*_ = 0 designates its absence. We also partition Ω into three disjoint groups^1^: (a) proteins Ω_0_ whose presence is externally stimulated by the given environment, (b) proteins 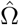 whose states influence the fitness value, and (c) the remaining dummy proteins Ω′.

A phenotype is then treated as a vector of zeros and ones, every of whose entries corresponds to the binary state of a protein in 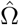. The resultant phenotype 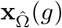 of a GRN *g* is determined by the reachability in *g*: For any 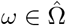, *x*_*ω*_ = 1 if and only if there is a stimulus *ω*_0_ ∈ Ω_0_ and a path from *ω*_0_ to *ω* in *g*, which represents a chain of sequentially triggered gene expression that generates protein *ω*. Finally, the fitness *f* is simply a function of the phenotype 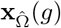.

Combined, a fitness landscape of GRNs is characterized through three key elements: the genotype network *G*, the external stimuli Ω_0_, and the fitness function of phenotype *f* (which implicitly identifies the fitness-relevant proteins 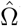). The genotype network *G* serves as the skeleton the fitness landscape, whereas the environment-dependent stimuli Ω_0_ and fitness function *f* determine the phenotypes of GRNs and their selective advantages, and they introduce definite fitness correlation between GRNs with the same accompanying phenotype.

## 3 Results

We present three theoretical insights of fitness landscape models using GRNs as the embedded genotype-phenotype mapping. First, we show that the according family of fitness landscapes must follow two topographical properties due to the high dimensionality of GRNs. Second, a mesoscopic skeleton of fitness landscapes is proposed, which is a more compact alternative to analyze evolutionary processes on fitness landscapes. Third, we provide a bottom-up approach to algorithmically construct this mesoscopic backbone of the fitness landscape of GRNs and demonstrate its better efficiency than coarse-graining the genotype network in a brute-force manner.

### 3.1 Connectivity and accessibility in a fitness landscape of GRNs

A fitness landscape model of GRNs features a handful of properties that have either been discovered in empirical fitness landscapes or investigated in the theoretical metaphor. On the one hand, its underlying space, i.e., the genotype network *G*, presents immense dimensionality since every locus of a GRN can potentially mutate into various expression behavior, which leads to a considerable amount of mutational neighbors. On the other hand, the fitness function *f* is flexible to tune the ruggedness of the fitness landscape. For example, a highly rugged holely landscape can be modeled by a binary *f* such that any GRN *g* ∈ 𝒢 has high fitness once some single protein 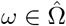 is present, *x*_*ω*_ = 1, and otherwise *g* becomes hardly fitted. Because one can always find several mutational neighbors of *g* whose phenotype shows an opposite state *x*_*ω*_, the resultant fitness landscape is inevitably rugged.

We further show that fitness landscape models of GRNs must hold the following two characteristics: First, for any two GRNs that share a common phenotype, there is a sequence of mutations connecting them through GRNs with the exactly same phenotype (with only one exception of an extreme phenotype). Second, for any two GRNs accompanying with two “not-so-distant” phenotypes (in terms of the number of proteins present in either one), there is always a mutational trajectory between them where the fitness is non-decreasing, regardless of the arbitrary fitness function. In other words, a fitness landscape of GRNs features both connectivity and accessibility.

Suppose that **y** is a phenotype. We use the notation 𝒢_**y**_ to denote the set of all GRNs with the resultant phenotype **y**, i.e., 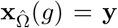 for *g* ∈ 𝒢_**y**_, under the given external stimuli Ω_0_. We also write 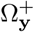 to be the proteins that are present in the phenotype **y**, so *x*_*ω*_ = 1 for 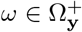 for and *x*_*ω*′_ = 0 for any other 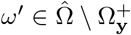. Note that the number of required-present proteins 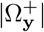 is bounded from above by the number of loci |Γ| since any present protein that is not an external must be triggered by the expression of some locus.

We observe that there are some GRNs 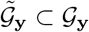 which play a “central” role among GRNs with the same phenotype **y**. Specifically, for any 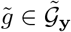, all the edges in 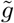 point from the stimuli Ω_0_ to the required-present proteins 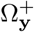, and each 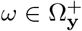 is targeted by at least one edge in 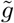. We demonstrate an example of such 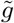 in Figure 1a. Crucially, these 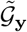are deemed central because they can be reached by any GRN *g* ∈ 𝒢_**y**_ through mutations among 𝒢_**y**_ themselves: First, for every edge in *g* that points to an 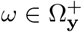, we rewire the edge such that it still points to *ω* but now from an *ω*_0_ ∈ Ω_0_. Arbitrarily rewiring the remaining edges between Ω_0_ and 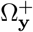 then leads to some central GRN in 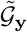 (see Figure 1a).

**Figure 1:**
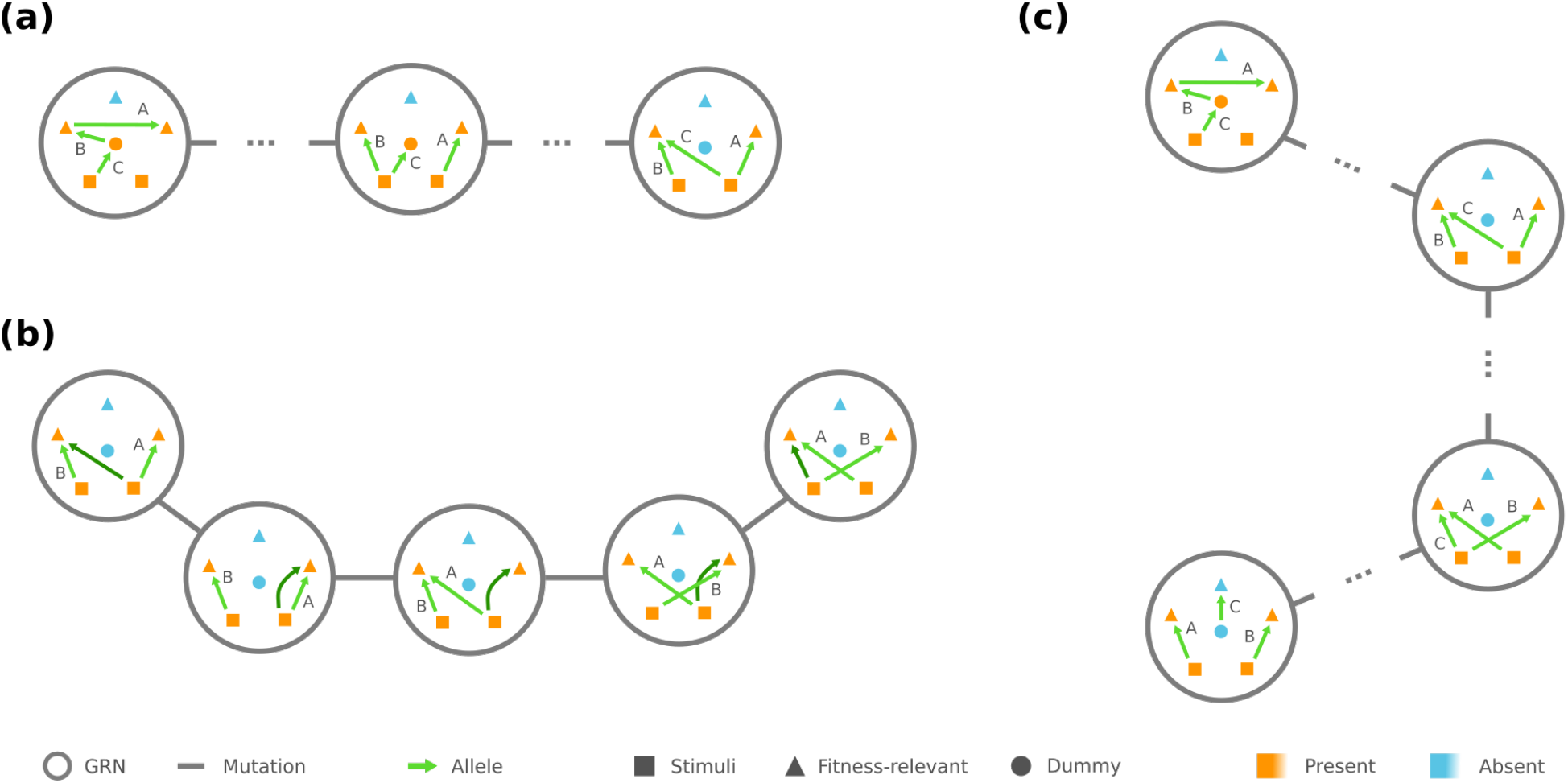
GRNs of the same phenotype are connected simply through themselves. (a) Any GRN can be rewired/ mutated into a “central” GRN (shown on the right). (b) A redundant edge (dark green) makes it feasible to turn any central GRN into another via edge rewiring, in this example exchanging the expression of locus A and B. (c) There is a mutational trajectory between any GRNS of the same phenotype through the central GRNs.

In addition, if the phenotype **y** has strictly less required-present proteins than the number of loci, the central GRNs 𝒢_**y**_ are mutually reachable by edge rewiring among 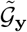: When 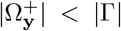, there is always a redundant edge^2^ whose rewiring makes no change in the phenotype. Such an edge helps us rewire each edge to any desired source/target pair between Ω_0_ and 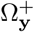 (see Figure 1b), which subsequently creates a chain of mutations between any 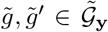. On the other hand, in the extreme case where 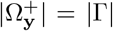, central GRNs 𝒢_**y**_ scatter over multiple connected components regarding mutations among 𝒢_**y**_ (detailed in Appendix A). These results implicate that, for any phenotype **y** with 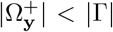 and any *g*_1_, *g*_2_ ∈ 𝒢_**y**_, there is always a mutational trajectory between *g*_1_ and *g*_2_ that only traverses over GRNs in 𝒢_**y**_, especially through the central ones 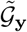 (see Figure 1c).

Next, we turn to mutations between two different phenotype **y** and **y**′. We observe that, if 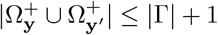, there are always two “peripheral” GRNs *ĝ* ∈ 𝒢_**y**_ and *ĝ*′ ∈ 𝒢_**y**′_ which only differ by one edge rewiring. To be more specific, there are two independent chains in *ĝ*, one of which begins with an stimulus *ω*_0_ ∈ Ω_0_ and sequentially connects the proteins required present in **y** but not in **y**′, i.e., 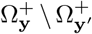, while the other consecutively joins 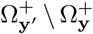. The rest of the edges in *ĝ* merely point from Ω_0_ to 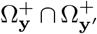, and each 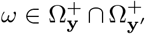 is targeted by at least one edge (see example in Figure 2, left). The other GRN *ĝ*′ only differ from *g* by the first edge in the chain of 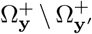, which is rewired such that it points from the stimulus *ω*_0_ to the first node in the chain of 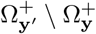 (Figure 2, right).

**Figure 2:**
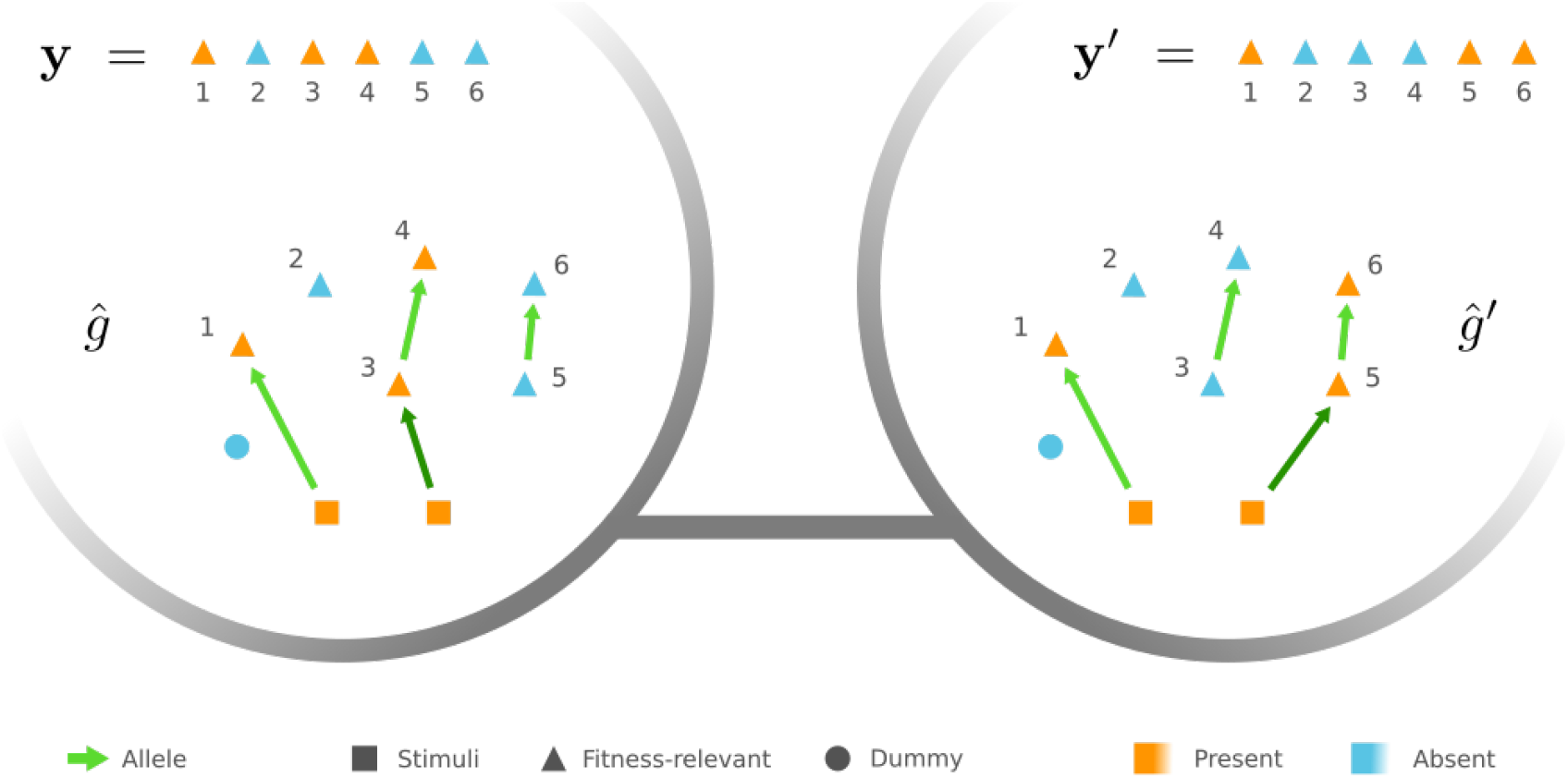
Example of peripheral GRNs connecting two different phenotypes. In the example peripheral GRN *ĝ* of phenotype **y**, there is a chain that triggers the presence state of proteins 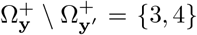. Contrarily, the other peripheral GRN *ĝ*′ of phenotype **y**′ contains a chain of proteins 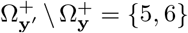. *ĝ* and *ĝ*′ are mutational neighbors since they only differ by rewiring the dark green edge, i.e., the first edge in either chain.

Since the GRNs 𝒢_**y**_ is mutationally connected for a phenotype **y** with 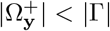, our observation suggests that there is a sequence of mutations from any GRN *g* ∈ 𝒢_**y**_ to a GRN *g*′ with an arbitrary phenotype **y**′ as long as 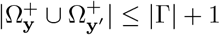. In particular, the mutational trajectory starting at *g* first traverses within 𝒢_**y**_ to a peripheral GRN and then transition into 𝒢_**y′**_ to reach *g*′, where the fitness is non-decreasing supposed that *f*(**y**′) ≥ *f*(**y**). Moreover, even under the extreme scenario 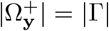, it is not hard to see that each central GRN in 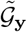can reach to an individual peripheral GRN through mutations among 𝒢_**y**_ (Appendix A), so the statement is also true for this extreme case. We also note that if the number of fitness-relevant proteins 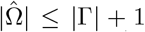, then the condition 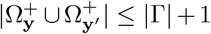 is assuredly satisfied for any two phenotypes **y** and **y**′. As a corollary, if 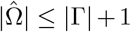, the fitness optimum will always be accessible.

To summarize, the genotype network *G* of GRNs can be imagined as the below metaphor: For a phenotype **y**, the subgraph of *G* induced by 𝒢_**y**_ organizes itself a single connected component due to the central GRNs 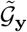, with the only exception that when 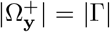, the induced subgraph shatters into multiple connected components. Furthermore, any two components of different phenotypes **y** and **y**′ with 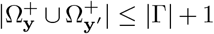 are connected through some peripheral GRNs. The genotype network *G* consequently ensures a fitter-and-fitter trajectory from between phenotypes that don’t largely differ in their required-present proteins.

### 3.2 Mesoscopic skeleton derived from “symmetries” in the genotype network of GRNs

Despite that the genotype network provides a great conceptualization of the inherent space of the fitness landscape, the number of all plausible GRNs grows even super-exponentially as the underlying loci and proteins expand. As a result, constructing the genotype network becomes extremely challenging beyond a small Γ and Ω. Here, we present a more compact skeleton of the fitness landscape of GRNs, where GRNs that play an equivalent role in the genotype network are consolidated through a “representative”, and GRNs in the same representative group feature the same phenotype.

As the underlying space of a fitness landscape of GRNs, the genotype network *G* is arguably a formulation with moderately redundant complexity. On the one hand, GRNs leading to the identical phenotype are deemed to have equal fitness. On the other hand, given any GRN, for example the mega-node rounded by orange in Figure 4a, one can always find some other GRN such that their neighborhoods in *G* are locally similar, e.g., the mega-node rounded by blue. In this example, we can easily see the similarity between the local neighborhoods of the two mega-nodes since the corresponding GRNs only differ by exchanging the role of locus A and B. This simple demonstration suggests that the structure of the genotype network *G* is not arbitrary; instead, some structural “symmeteries” exist.

In graph theory, symmetries in a network are formally described through its automorphisms. An automorphism of a graph is a way to shuffle the labels of its nodes such that the graph remains identical before and after shuffling. For instance, in Figure 3b, exchanging node 2 and 3 generates the same network and is thus an automorphism, whereas exchanging node 2 and 4 is not because there is an edge from 2 to 3 after shuffling, but such an edge doesn’t exist before shuffling. To be more concrete, an automorphism of the genotype network *G* is a permutation^3^ *σ* of all plausible GRNs 𝒢 = *V*(*G*) such that, for any *g*_1_, *g*_2_, ∈ 𝒢 (*σ*(*g*_1_), *σ*(*g*_2_)) ∈ *E*(*G*) if and only if we also have (*g*_1_, *g*_2_) ∈ *E*(*G*). Importantly, once two GRNs *g* and *g*′ are related through an automorphism *σ* of *G*, e.g., *g*′ = *σ*(*g*), it is not hard to envision them playing an equivalent role in *G*. Specifically, these two GRNs share the same mega-node properties as long as the properties are fully determined by the connections in the genotype network (see Proposition B.1).

**Figure 3:**
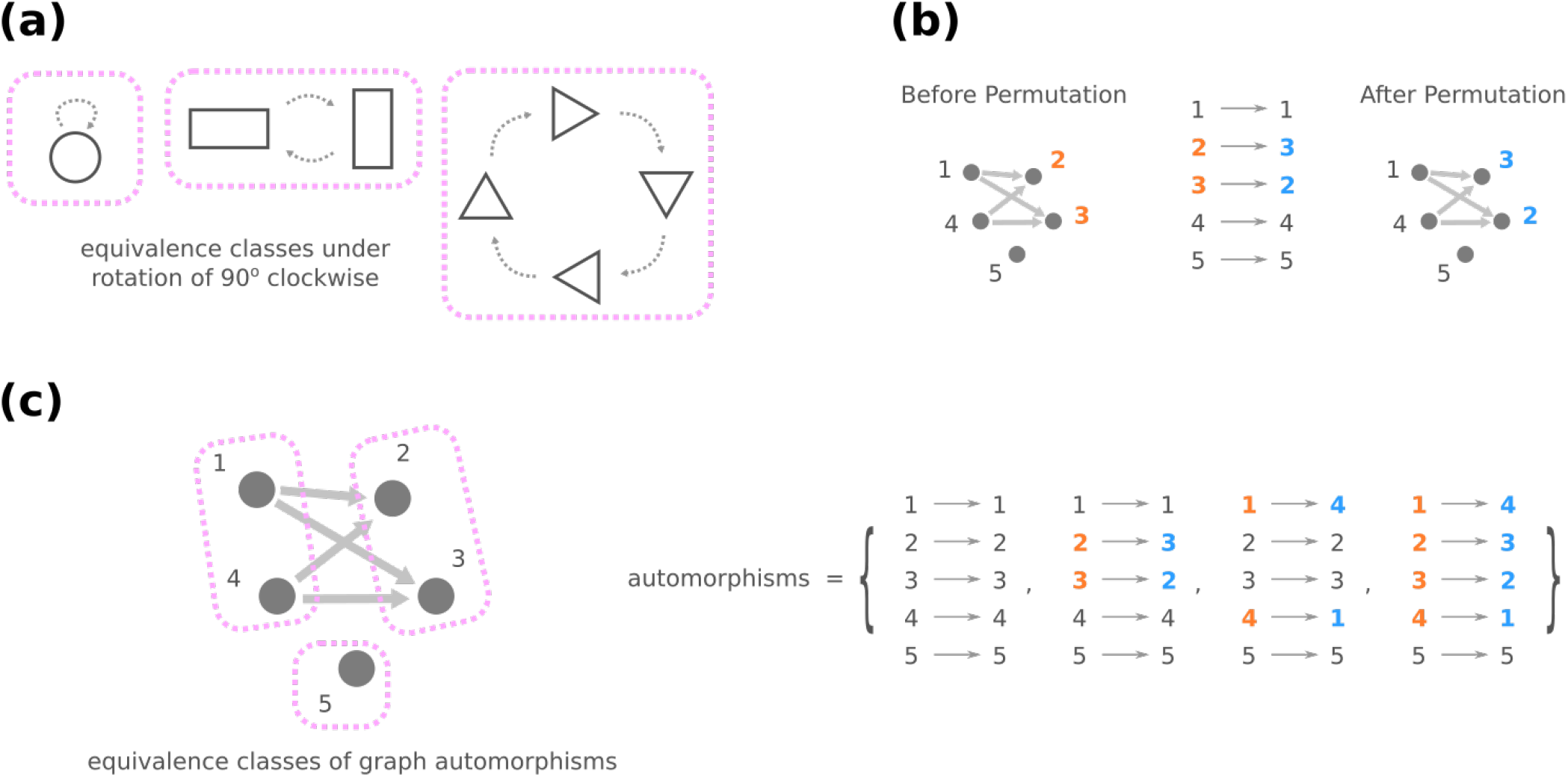
(a) As a minimal example, let’s imagine an operation that rotates a geometric object by 90 degrees clockwise. The rotation maps one object into another (dashed arrows), and it leads to equivalence classes where objects are grouped by their symmetry under rotation (pink rectangles). (b) An automorphism of a graph is a permutation of nodes that retains the same graph. (c) Equivalence classes under graph automorphisms bring together nodes that have similar roles connection-wise in the graph.

Furthermore, automorphisms partition the GRNs by their roles in the genotype network through the mathematical concept of equivalence classes. For a high-level and general description, imagine a set of elements and a group of operations acting on them. Each operation turns one element into another, and these two elements are related by the operation to indicate some similarity between them. An equivalence class consists of elements that are mutually related by any operation, and the set of elements are said to be partitioned into equivalence classes under the action of the operations (see Figure 3a for an illustrative example). Going back to automorphisms of a graph *G*′, if the group of all automorphisms Σ(*G*′) is known, the equivalence classes of nodes *V*(*G*′) under the action of Σ(*G*′) then gather nodes with a similar “structural position” in *G*′ (Figure 3c).

In order to find a more compressed, less redundant backbone of a fitness landscape of GRNs, we are not only interested in automorphisms that relate GRNs by their similar roles in the genotype network, but these automorphisms also need to preserve the phenotype after mapping one GRN to another. We use the notation Σ_*x*_(*G*) for the set of such automorphisms of *G*, i.e., for any *σ* ∈ Σ_*x*_(*G*) and GRN *g* ∈ 𝒢, *σ*(*g*) and *g* have the same phenotype. The equivalence classes of mega-nodes *V*(*G*) under the action of phenotype-preserving automorphisms Σ_*x*_(*G*) then unite GRNs that (a) show similar mutational relationship with others and (b) lead to the same fitness due to their identical phenotype. We will mildly abuse the terminology to call them the *equivalence classes of GRNs*, which we denote by Θ and each *θ* ∈ Θ is a set of GRNs related through Σ_*x*_(*G*). Crucially, since the mutational relationship and the resultant phenotype are the two components that characterize a GRN in the fitness landscape, GRNs in a *θ* ∈ Θ are deemed equivalent (semantically beyond the mathematical terminology) and they can be reduced to an arbitrary representative among them. Therefore, the equivalence classes of GRNs provide an efficient way to depict the underlying space of the fitness landscape.

Yet, what exactly are the phenotype-preserving automorphisms Σ_*x*_(*G*) of the genotype network? From a sufficiency direction, we show that there exist a few graphical operations on the GRNs that produce phenotype-preserving automorphisms. These graphical operations involve permuting/shuffling different sorts of elements in a GRN:

i. The identities of loci Γ, e.g., exchanging edge labels of locus A and B in Figure 4b.
ii. The identities of dummy proteins Ω′, e.g., exchanging node labels of protein 3 and 4 in Figure 4c. and
iii. Change the source node of an edge from one stimulus to another stimulus and vice versa, e.g., in Figure 4d, moving an edge pointing from node 1 to node 3 to point from node 2.
iv. Move a self-loop at one node to another node and vice versa, for example, re-allocating a self-loop at node 3 to node 4 in Figure 4e.

**Figure 4:**
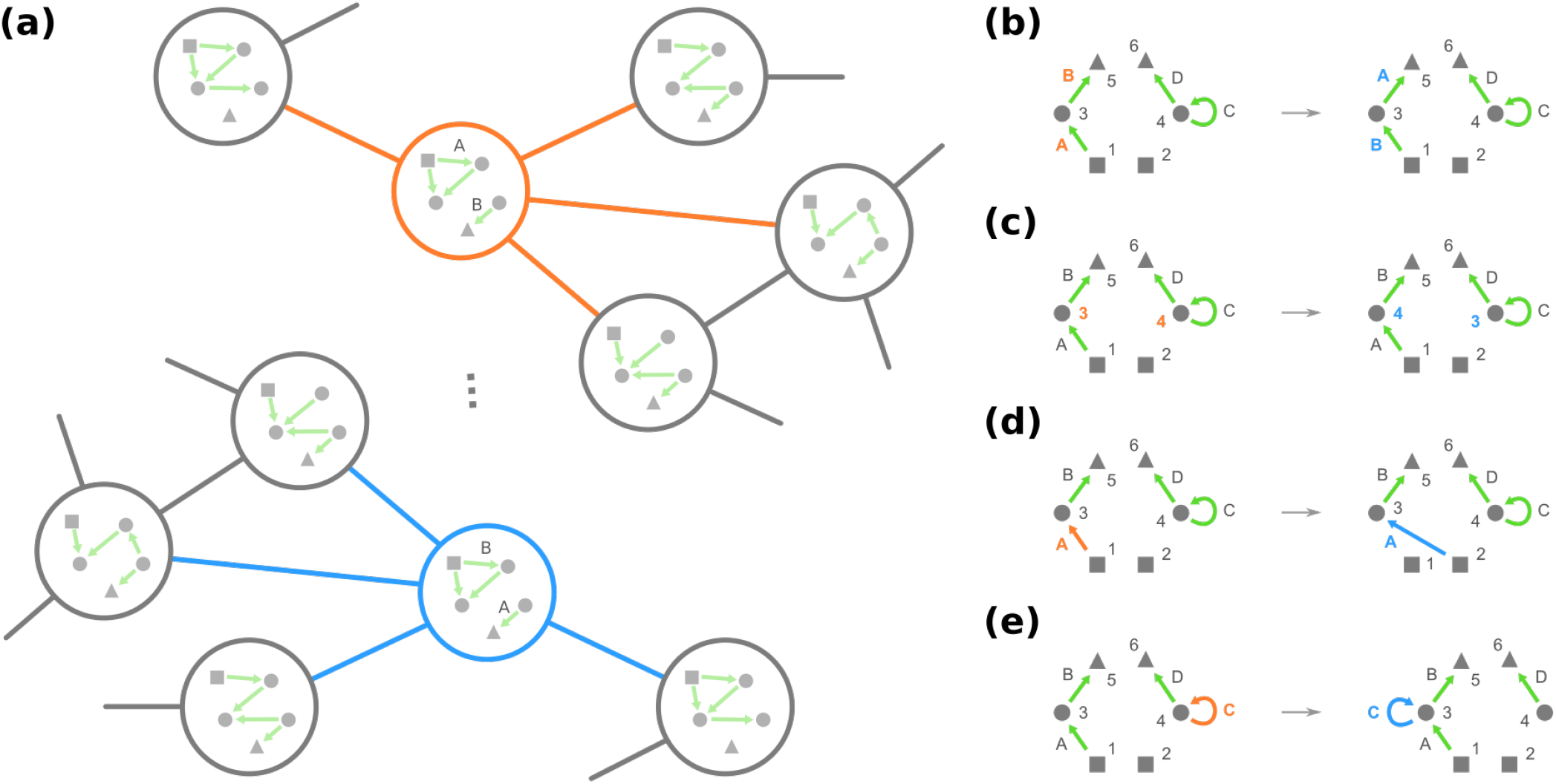
The genotype network has some symmetry such that multiple GRNs have similar local neighborhoods, as we demonstrate in (a). More formally, these GRNs constitute an equivalence classes under phenotype-preserving automorphisms, which can be found by graphical operations of (b) permuting loci, (c) permuting dummy proteins (circles), (d) exchanging edges pointing from two different stimuli (squares), if any, and (e) exchanging self-loops at two different nodes, if any.

We leave the formal proofs in Theorem B.1, where the idea is that these graphical operations preserve the phenotype of GRNs and facilitate mapping one mutational connection to another in the genotype network *G*. Note that combining multiple operations of category (iii) and (iv) implicates that if two GRNs can be transformed from one into the other via (a) rewiring edges as long as they remain to point from stimuli or (b) arbitrarily allocating self-loops, then they must belong to the same equivalence class.

On the other hand, from a necessity direction, one can computationally obtain a partition^4^ 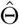 of the GRNs 𝒢 that is coarser^5^ than the equivalence classes Θ. Specifically, start with a partition 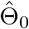 where GRNs with the same resultant phenotype are grouped together. We create a sequence of partitions of 𝒢 through the following iterative procedure: Given the partition 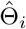, the next partition 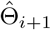 is obtained by further dividing groups in 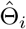 (if needed) such that for each group 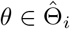 and 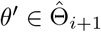, any two GRNs in *θ*′ have the same number of neighbors among *θ* in the genotype network *G*. This iterative procedure is terminated when no further division is required, i.e., 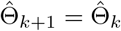 for some integer *k* (see Figure 5a for an illustrative cartoon of the iterative procedure). We then have 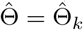 to be our desired partition of GRNs.

**Figure 5:**
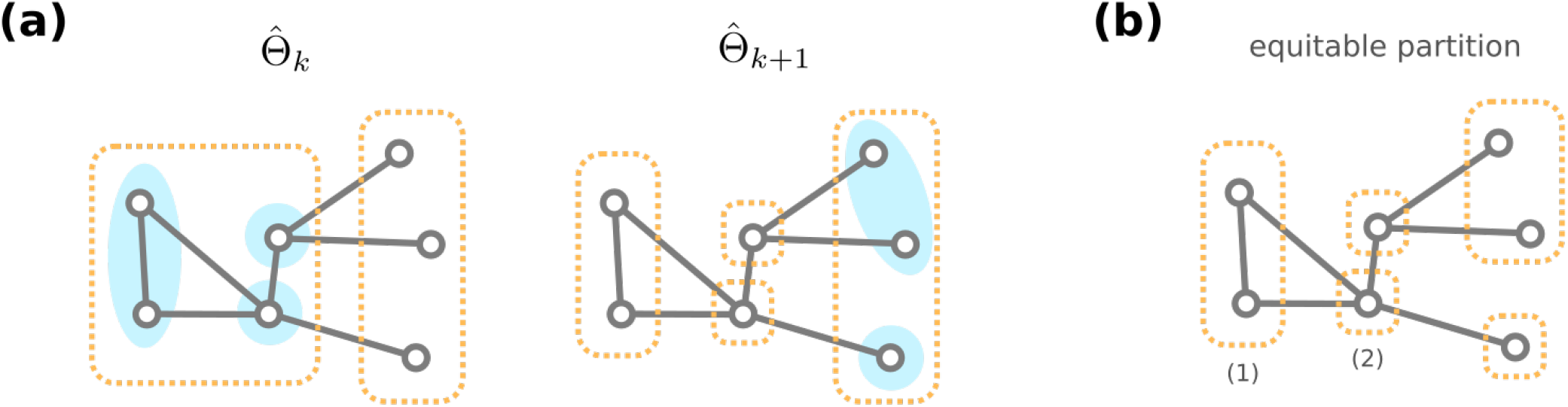
(a) Consider a fake, toy example genotype network of GRNs (here we omit the exact content of GRNs). Given the partition 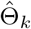, note that mega-nodes in a group (dashed orange rectangle) may share different number of connections among other groups (blue shaded circles), and they are further divided to generate the next partition 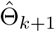. (b) Both the equivalences classes of GRNs and the stationary partition from our iterative procedure are equitable, e.g., either of the two mega-nodes in group (1) has one connection among (1), another connection with (2), and none with other groups.

To see why the proposed iterative procedure generates a coarser partition 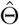 than the equivalence classes Θ of GRNs, we stress that the equivalence classes under the action of a group of automorphisms always form an equitable partition of the nodes. A partition 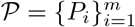 of nodes of a graph is equitable [68] if for every *P*_*i*_, *P*_*j*_ ∈ 𝒫, any two nodes *u, v* in group *P*_*i*_ have the same number of neighbors in *P*_*j*_ (Figure 5b). Since the equivalence classes under automorphisms gather nodes that share a similar structural role/position in the graph, and such similarity includes how many neighbors they have in other equivalence classes, they are expected to be an equitable partition. Combined with the fact that GRNs in an equivalence class *θ* ∈ Θ must have the same amount of neighbors for each different phenotype, we inductively show that any two GRNs *g*_1_, *g*_2_ ∈ *θ* are never separated during the iterative procedure that generates 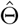 (see Theorem B.2). Therefore, any equivalence class *θ* ∈ Θ must be included in a computationally acquired group 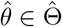.

Figure 6 demonstrates the coarser partition 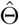 generated by the iterative procedure for an arbitrary toy example. The obtained 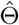 contains 154 groups of GRNs, and the size of groups ranges from 2 to 96. We also count the number of different kinds of GRNs that can not be mutually transformed through graphical operations (i) and (ii), and this number varies from 1 to 4 in our example 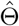. Moreover, for every group in 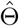, we observe that those different kinds of GRNs can be transformed from one to another by changing the stimulus an edge is pointing from and re-allocating self-loops (e.g., see Figure 6b). 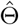 is thus not simply a coarser partition than the equivalence classes; according to (i)–(iv), we know that groups in 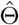 are exactly the equivalence classes Θ. This arguably general enough toy example implicates that there is no need of other graphical operations to determine the equivalence classes of GRNs.

**Figure 6:**
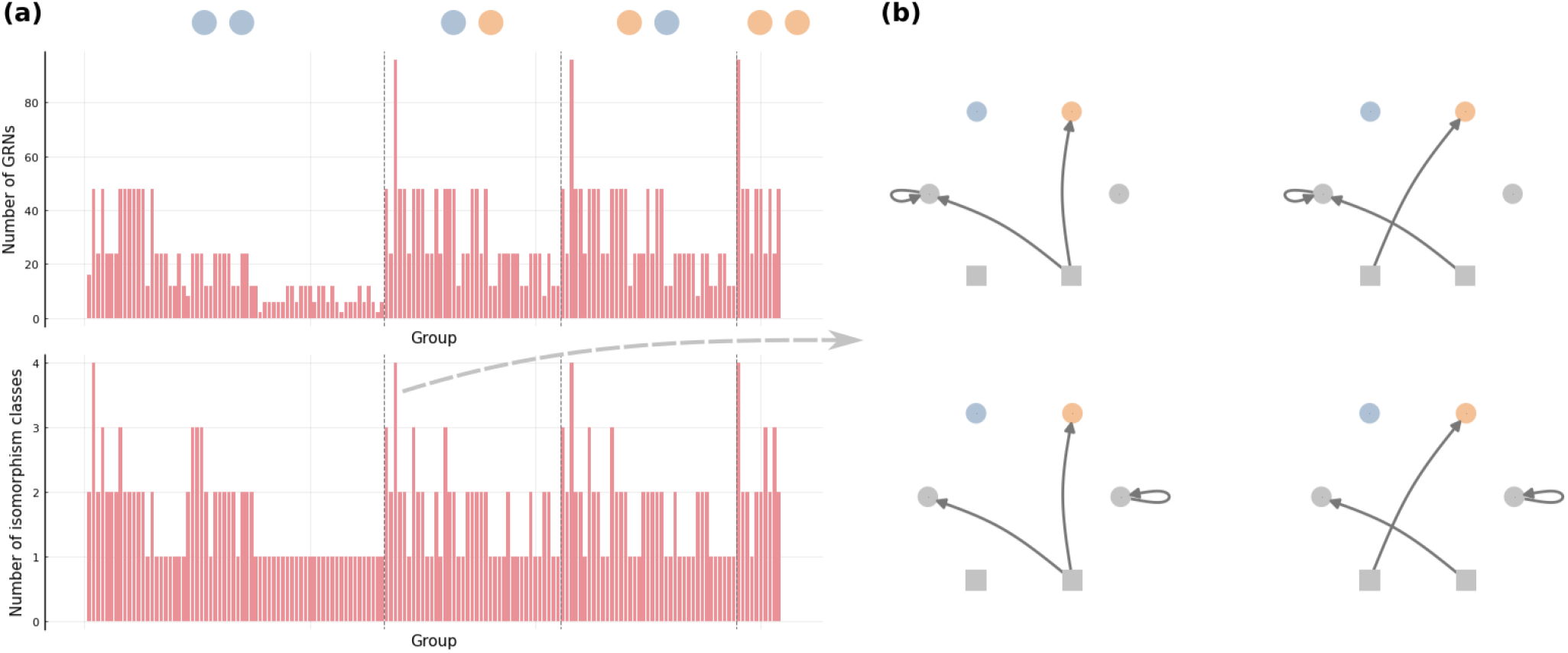
Example partition coarser than the equivalences classes of GRNs. We run the proposed iterative procedure with |Γ| = 3 and 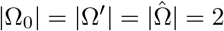, where stimuli Ω_0_ are drawn as squares and the present/absent state of fitness-relevant proteins 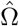 are colored by orange/blue. (a) The number of GRNs and the number of isomorphism classes of GRNs in each group of the obtained partition 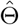, where the dashed lines separate groups of different phenotypes. (b) Isomorphism classes of GRNs in a group.

As a result, we conjecture that, in general, all the phenotype-preserving automorphisms Σ_*x*_(*G*) of the genotype network can be generated through combining graphical operations (i) to (iv) on the GRNs. More importantly, we now understand what makes two GRNs *g*_1_ and *g*_2_ in the same equivalence class *θ* ∈ Θ: After removing all the self-loops merging stimuli Ω_0_ into a single node, there are permutations of loci Γ and dummy proteins Ω′ that jointly transform *g*_1_ into *g*_2_. This condition reconciles with the concept of isomorphisms between graphs. Whereas as an automorphism is a mapping of nodes such that a graph preserves itself, an isomorphism is a mapping of nodes that transform one graph into another (and its definition can be easily extended for graphs with edge labeling as well). We will borrow the terminology and call the two permutations of Γ and Ω′ together a *phenotype-preserving isomorphism* from *g*_1_ to *g*_2_. Therefore, in other words, two GRNs belong to the same equivalence class if and only if there is a phenotype-preserving isomorphism between them after self-loop removal and stimuli merging.

### 3.3 Algorithmic construction of the mesoscopic backbone of GRN fitness landscape

Next, we turn to algorithmic approaches which establish the underlying space of a fitness landscape of GRNs, where our goal is to investigate any possibility that is more computationally efficient than the brute-force construction of the genotype network *G*. Since the equivalence classes Θ coarse-grain GRNs with the same phenotype and a similar structural role in *G*, and a representative GRN is adequate to entail all other GRNs in an equivalence class, we anticipate an effective algorithm to build up the underlying space of the fitness landscape through these equivalence classes of GRNs. In particular, a desired algorithm must (a) acquire the equivalence classes Θ from scratch, and (b) for a representative GRN in any equivalence class, count the number of its mutational neighbors in other equivalence classes and also within the class it belongs to.

To avoid any confusion, we emphasize that, although the task of drawing mutational connections between equivalence classes Θ is seemingly simply grouping mega-nodes in the genotype network *G*, this naive network science exercise is an unsuitable (and even contradictory) approach in our scenario. First and for most, it is the heavy computational expense of constructing *G* that motivates our search on a more compact backbone of the fitness landscape of GRNs. Yet, grouping mega-nodes in *G* demands prior knowledge of the genotype network *G* itself. Second, in contrast to coarse-graining nodes in a graph where the groups of nodes are pre-specified, listing all GRNs in an equivalence class requires examining pairs of GRNs and assuring a phenotype-preserving isomorphism between them after removing self-loops and merging stimuli. Determining the equivalence classes Θ from all the GRNs 𝒢 = *V*(*G*) can thus be as costly as building the genotype network *G*. These two reasons again resonate the value of the equivalence classes Θ, which consolidate GRNs into their equivalent representatives.

Here, we present a bottom-up approach that enumerates each equivalence class of GRNs and simultaneously compute the number of mutational connections among them. To begin, recall from section 2.2 that mutation from a GRN *g*_1_ ∈ 𝒢 to another *g*_2_ ∈ 𝒢 corresponds to rewiring a single edge in *g*_1_. The edge rewiring can be further designated into one of the four scenarios: *g*_1_ may rewire a self-loop/non-self-loop edge to a self-loop/non-self-loop edge in *g*_2_. We observe that if *g*_1_ and *g*_2_ are mutational neighbors, then the number of non-self-loop edges in *g*_1_ and *g*_2_ differ at most by one. Notation-wise, we will write Γ′(*g*) as the loci representing the non-self-loop edges in the GRN *g*, and |Γ′(*g*)| as the number of those non-self-loop edges. In other words, given an equivalence class *θ* ∈ Θ and a representative GRN *g* ∈ *θ*, there is undoubtedly no mutational neighbor of *g* in another equivalence class *θ*′ (with a representative GRN *g*′ ∈ *θ*′) where the difference between |Γ′(*g*)| and |Γ′(*g*′)| is greater than one.

We can therefore establish the underlying space of the fitness landscape by incrementally examining each equivalence class with a increasing number of non-self-loop edges in the representative GRN. This strategy is envisioned in Figure 7, where the underlying space can be viewed as “layers” of equivalence classes of GRNs. Let Θ_*k*_ be the set of equivalence classes where for every *θ* ∈ Θ_*k*_, the representative GRN *g* ∈ *θ* has exactly *k* non-self-loop edges, |Γ′(*g*)| = *k*. We start with the layer Θ_0_, which consists of the only equivalence class with no non-self-loop edges. Then, with layers Θ_0_, Θ_1_, …, Θ_*k*_ and all the mutational connections among them, we will find the equivalence classes in the next layer Θ_*k*+1_ and their mutational connections with layer Θ_*k*_ and within themselves, up until *k* = |Γ| where all edges are not self-loops.

**Figure 7:**
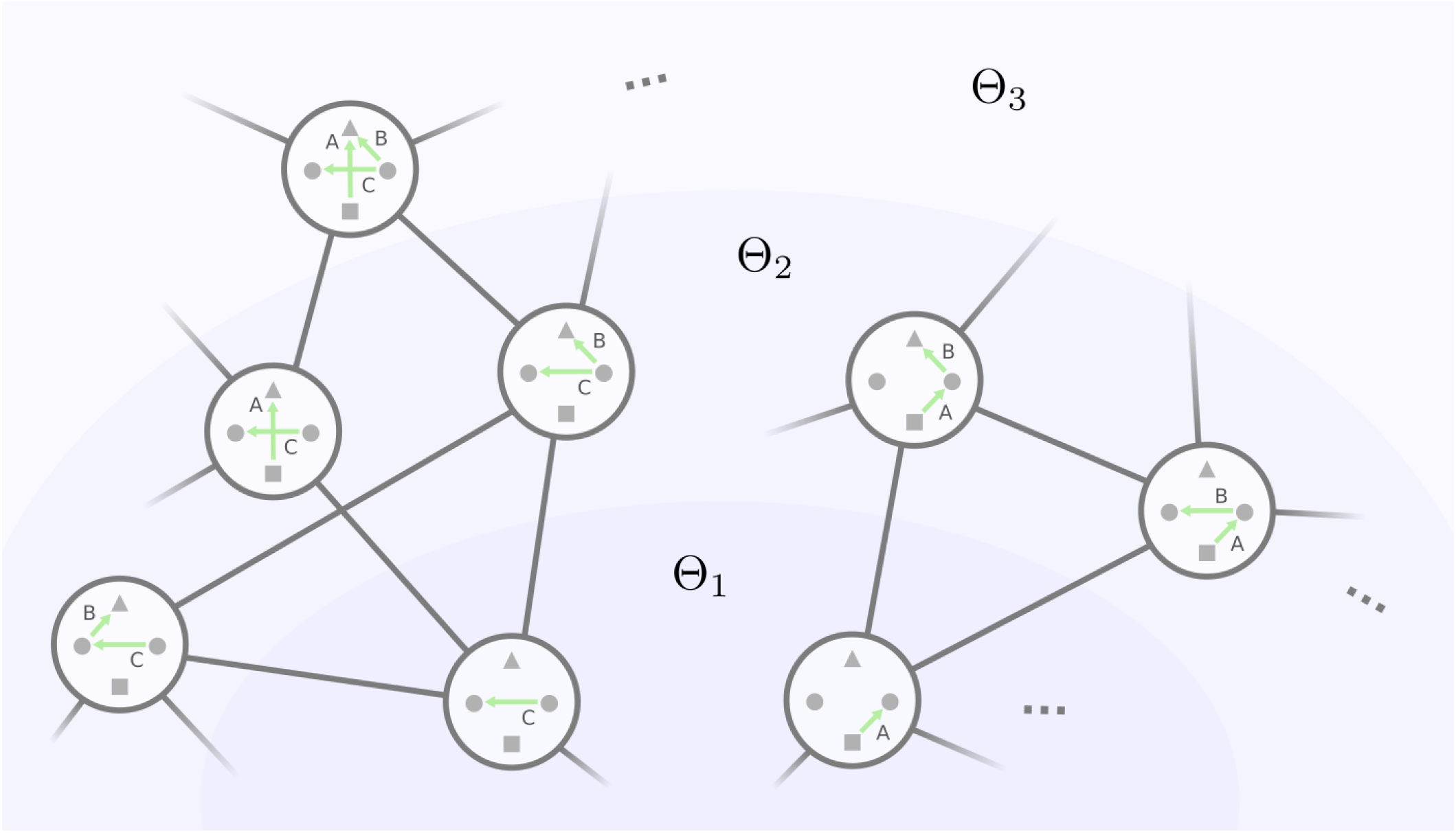
Layering the GRNs by their number of non-self-loop edges. A GRN’s mutational neighbors must fall into the same or the adjacent layers. For the ease of illustration, we only show the non-self-loop edges neglect the protein states in GRNs.

To be more precise, for any GRN *g* ∈ 𝒢, we use the notation ℳ^+^(*g*) to denote the mutational neighbors of *g* that have one more non-self-loop edge than *g*. This notion of mutational neighbors with one more non-self-loop edge, which we abbreviate as the ℳ^+^ neighborhood, is sufficient to capture the relationship between *g* and its arbitrary mutational neighbor *g*′:

- If *g*′ has one more non-self-loop edge than *g*, then *g*′ ∈ ℳ^+^(*g*).
- If *g*′ has one less non-self-loop edge than *g*, then we have *g* ∈ ℳ^+^(*g*′).
- If *g*′ has the same number of non-self-loop edge as *g*, then they share a common mutational neighbor *g*″ where the only different edge between *g* and *g*′ is rewired to a self-loop, and thus *g, g*′ ∈ ℳ^+^(*g*″).

The mutational connections between equivalence classes can hence be uncovered through investigating the ℳ^+^neighborhood of the representative GRN for each equivalence class. Moreover, the ℳ^+^neighborhood of representative GRNs in layer Θ_*k*_ reveal the equivalence classes in layer Θ_*k*+1_ because any GRN must have a mutational neighbor with one less non-self-loop edge. However, the question is how to join different ℳ^+^ neighbors into equivalence classes. In particular:

A. For an equivalence class *θ* ∈ Θ_*k*_ and its representative GRN *g* ∈ *θ*, under what condition will 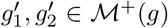 belong to the same equivalence class in layer Θ_*k*+1_?
B. For two distinct equivalence classes *θ*_1_, *θ*_2_ ∈ Θ_*k*_ and their representative GRNs *g*_1_ ∈ *θ*_1_ and *g*_2_ ∈ *θ*_2_, under what condition will 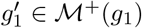 and 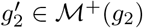 belong to the same equivalence class in layer Θ_*k*+1_?

For our ease to exempt from the stimuli merging action, we hereafter choose the GRNs *g, g*_1_, *g*_2_, 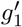 and 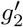 such that only one stimulus node is incident to out-going edges.

To address (A), we first suppose that 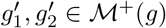 belong to the same equivalence class, so there is a phenotypepreserving isomorphism *π* from 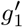 to 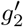 after self-loop removal. For the ease of demonstration, here we abuse this notation *π* to collectively represent the permutations^6^ over the loci Γ and dummy proteins Ω′. We also recall from section 2.2 that we write *e*_*g*_(*γ*) = (*u, v*) to denote “the source-target pair of edge *γ* is (*u, v*) in GRN *g*.” Furthermore, we write 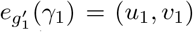 and 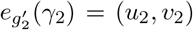 where *γ*_1_ and *γ*_2_ are the edge added to *g* that forms 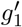 and 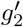 respectively (addition in terms of rewiring a self-loop to a non-self-loop edge). A few observations follow assuming such an isomorphism *π* exists:

1. There is an integer *p* such that *π*^*p*^(*γ*_1_) = *γ*_1_ and (*π*^*p*^(*u*_1_), *π*^*p*^(*v*_1_)) = (*u*_1_, *v*_1_).
2. There is another integer *q* < *p* such that *π*^*q*^(*γ*_1_) = *γ*_2_ and (*π*^*q*^(*u*_1_), *π*^*q*^(*v*_1_)) = (*u*_2_, *v*_2_).
3. 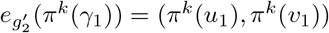 for *k* = 1, 2, …, *q*.
4. 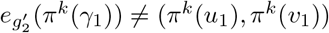 for *k* = *q* + 1, *q* + 2, …, *p*.
5. For any locus *γ* and non-self-loop source-target pair (*u, v*) with no 0 ≤ *k* ≤ *q* − 1 such that (*γ, u, v*) = (*π*^*k*^(*γ*), *π*^*k*^(*u*_1_), *π*^*k*^(*v*_1_)), we have *e*_*g*_(*π*(*γ*)) = (*π*(*u*), *π*(*v*)) if and only if *e*_*g*_(*γ*) = (*u, v*).

We detail the reasoning behind these observations in Lemma C.1–C.3. Critically, our fifth observation implies that, after self-loop removal, the isomorphism *π* between 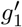 and 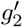 is in fact a phenotype-preserving automorphism of a subgraph 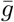 of the GRN *g*. In addition, observation 3. and 4. depict that those edges in *g* but not in 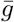 are sequentially mapped from one to another via this automorphism *π*, i.e., 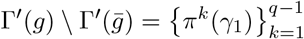, and they bridge the newly added edges *γ*_1_ and *γ*_2_ = *π*^*q*^(*γ*_1_) (along with the corresponding source-target pairs). We show that the converse is also true (see Theorem C.1): If we find a phenotype-preserving automorphism *π* of a subgraph 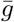 of *g* where *γ*_1_ is consecutively mapped to *γ*_2_ through the edge differences after self-loop removal 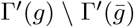, *π* is guaranteed a phenotype-preserving isomorphism from 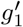to 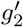 after removing the self-loops.

We now know the sufficient and necessary condition for two ℳ^+^ neighbors of *g* to be in the same equivalence class. Intriguingly, the key lies upon the phenotype-preserving automorphisms of subgraphs of the representative GRN *g*. Here we demonstrate a few simple examples in Figure 8a: In the top row, an automorphism of *g* directly maps between the two additional edges (*u*_1_, *v*_1_) = (3, 5) and (*u*_2_, *v*_2_) = (1, 4). In the middle row, the two edges (*u*_1_, *v*_1_) = (1, 2) is consecutively mapped to (*u*_2_, *v*_2_) = (3, 4) through the edge (2, 3), and (*u*_2_, *v*_2_) is consecutively mapped back to (*u*_1_, *v*_1_) through the non-edge (4, 1), so we have *q* = 2 and *p* = 4. As a mixture of both, in the bottom row, (*u*_1_, *v*_1_) = (2, 5) is consecutively mapped to (*u*_2_, *v*_2_) = (3, 6) through the edge (1, 4), and this isomorphism is exactly an automorphisms of a subgraph 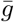 of *g* where the edge (1, 4) is removed.

**Figure 8:**
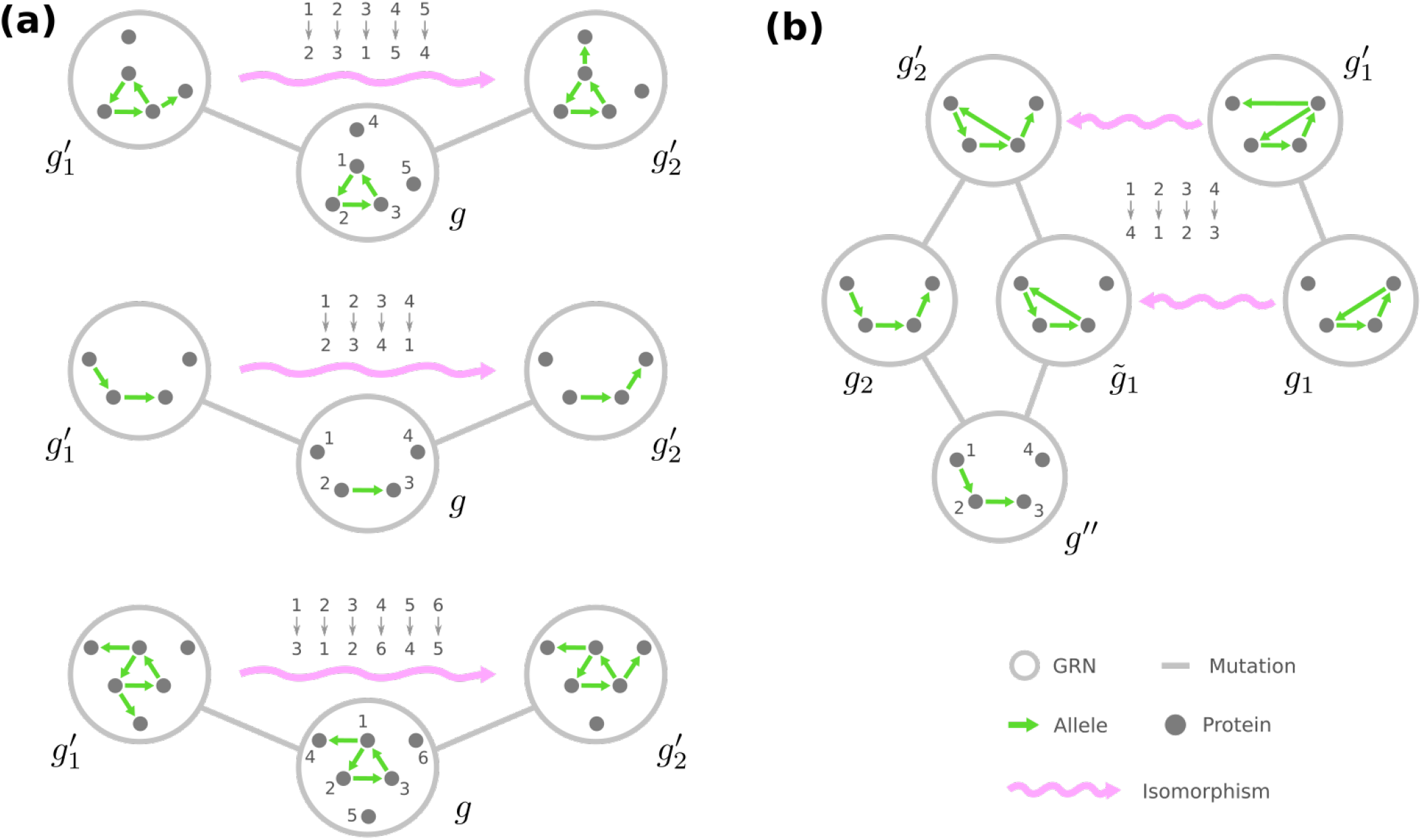
Sufficient conditions that two ℳ^+^ neighbors belong to an equivalence class. For the illustration purpose, we only show the dummy proteins and omit the protein states, edge labels and self-loops in (a) three examples such that two ℳ^+^ neighbors of a GRN *g* are isomorphic, and (b) an example where the ℳ^+^ neighbors 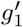 and 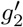 of GRNs *g*_1_ and *g*_2_ in different equivalence classes are isomorphic.

Let’s switch gear to tackle the remaining question (B). Suppose that *g*_1_ and *g*_2_ are the representative GRN in two different equivalence classes such that |Γ′(*g*_1_)| = |Γ′(*g*_2_)|. We in a similar vein start with the scenario where 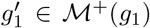 and 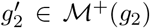 belong to the same equivalence class. Let *γ*_1_ and *γ*_2_ be the newly added edges to *g*_1_ and *g*_2_ that generate 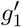 and 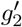 respectively, where 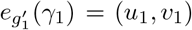 and 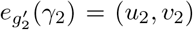, and let *π* be a phenotype-preserving isomorphism from 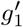 to 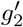 after self-loop removal. We observe that applying the permutation *π* on *g*_1_ transforms it into another GRN 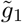 in the same equivalence class. Since *g*_1_ simply has one less edge *γ*_1_ than 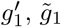 and 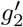 only differ by a missing edge *π*(*γ*_1_). Namely, we have 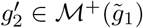 where, compared to 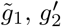 has the additional edge with 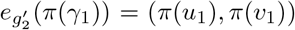. Moreover, remind that 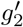 also belongs to the ℳ^+^neighborhood of *g*_2_ with the additional edge 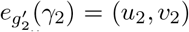. As a result, by removing both the extra edges from 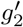, we find a GRN *g*″ such that 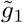, *g*_2_ ∈ ℳ^+^(*g*″).

We again present an illustrative example in Figure 8b. Here a GRN 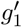 in the equivalence class of *g*_1_ can be found via the isomorphism *π* between 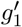 and 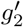. We note that the newly added edge (*u*_1_, *v*_1_) = (4, 1) is transformed to (*π*(*u*_1_), *π*(*v*_1_)) = (3, 4) in 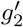, which is missing in 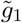. Removing both (*π*(*u*_1_), *π*(*v*_1_)) = (3, 4) and (*u*_2_, *v*_2_) = (3, 1) from 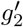 produces a GRN *g*″ which is a common neighbor of *g*_2_ and 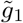 with one less non-self-loop edge.

Our observation resolves the necessary condition of (B): For the representative GRN of two different equivalence classes *g*_1_ ∈ *θ*_1_ and *g*_2_ ∈ *θ*_2_, if their ℳ^+^neighbor 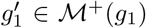 and 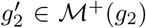 belong to the same equivalence class, then we can always find two GRNs 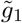 and *g*″ such that (a) 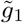 falls in the equivalence class of *g*_1_, and (b) 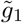 and *g*_2_ are ℳ^+^ neighbors of *g*″. Moreover, it is not hard to see that the converse is true as well (Theorem C.2). Therefore, in a nutshell, whether the ℳ^+^ neighborhood of *g*_1_ and *g*_2_ reveal a common equivalence class depends on the existence of a GRN *g*″ that the equivalence classes *θ*_1_ and *θ*_2_ are both rooted from.

With the resolutions of question (A) and (B) which implicate how to collect the ℳ^+^ neighbors of representative GRNs into equivalence classes, we now describe our algorithm that incrementally generates the equivalences classes Θ of GRNs and establishes the mutational connections among them. Suppose that we have already built layers of equivalence classes Θ_0_, Θ_1_, …, Θ_*k*_ and determined the mutational connections among them. For each representative GRN *g* in layer Θ_*k*_ and every *g*′ ∈ ℳ^+^(*g*), we will view *g*′ as the combination of *g* and an additional, non-self-loop edge with label *γ* ∈ Γ that points from *u* ∈ Ω to *v* ∈ Ω, for which we write *g*′ = *g* ⊕ (*γ, u, v*). All such plausible combinations lead to a collection of ℳ^+^neighbors of the representative GRNs in layer Θ_*k*_, for which we abuse the notation ℳ^+^(Θ_*k*_).

We initially put each *g*′ ∈ ℳ^+^(Θ_*k*_) into an individual group, and we define a collection of operations Φ that join groups of ℳ^+^neighbors. The operations Φ are of three different types:

I. For every representative GRN *g* in Θ_*k*_ and every phenotype-preserving automorphism *σ* of *g*, there is an operation *ψ*_*g,σ*_ that joins together the groups of 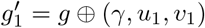 and 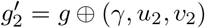 .where *u*_1_, *u*_2_ ∈ Ω_0_ and *v*_2_ = *σ*(*v*_1_).
II. For every representative GRN *g* in Θ_*k*_ and every phenotype-preserving automorphism 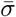 of each subgraph 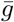 of *g* such that the edge differences 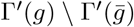 are sequentially connected via 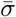 (along with their source-target pairs), there is an operation 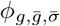 that joins together the groups of 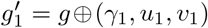 and 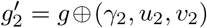, where automorphism 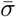 consecutively transforms edge *γ*_1_ into *γ*_2_ through 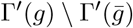.
III. For every representative GRN *g*″ in Θ_*k*−1_ and each 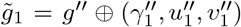 and 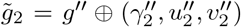 in two different equivalence classes *θ*_1_ and *θ*_2_, such that we have the phenotype-preserving isomorphism *π*_1_/*π*_2_ from 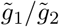 to the representative GRN *g*_1_/*g*_2_ after self-loop removal, there is an operation 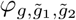 that joins together the groups of 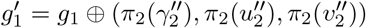 and 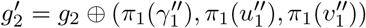.

The consequent groups of after applying the joining operations Φ then constitute the equivalence classes in the next layer Θ_*k*+1_. We hereafter write 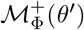 to denote the corresponding consequent group of an equivalence class *θ*′∈ Θ_*k*+1_. More importantly, we can then choose an arbitrary ℳ^+^ neighbor in 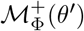 as the representative GRN of the equivalence class *θ*′, such that only one stimulus node is incident to out-going edges in the chosen representative GRN.

The joining operations Φ further provides useful information to count the number of mutation neighbors that a representative GRN *g* ∈ *θ* in layer Θ_*k*_ has among any equivalence class *θ*′, which we will denote by *A*_*g*_(*θ*′). Let’s first consider the scenario that *θ*′∈ Θ_*k*+1_. For any GRN 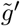 in the corresponding Φ-coupled group 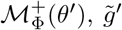 is a mutational neighbor of *g* if it can be viewed as a combination of *g* and an arbitrary extra non-self-loop edge, e.g., 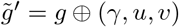 for some edge 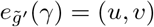. Hence, we have

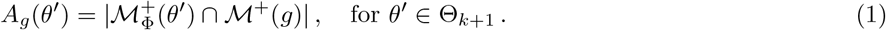

Note that in this case *A*_*g*_(*θ*′) is easily acquired when building up the layer Θ_*k*+1_ through Φ.

Second, for *θ*′ ∈ Θ_*k*−1_, *A*_*g*_(*θ*′) can be computed given *A*_*g*′_(*θ*), where *g*′ is the representative GRN of *θ*′. Due to the fact that the equivalence classes Θ generate an equitable partition of the genotype network *G* (see section 3.2), we have *A*_*g*_(*θ*′) × |*θ*| = *A*_*g*′_ (*θ*) × |*θ* ′ | equal to the total number of mutational connections between *θ* and *θ*′. Moreover, we can compute the size of the equivalence class *θ* via (see Appendix D)

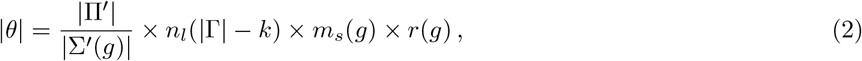

where (a) we use Π′ to denote the set of all permutations dummy proteins Ω′ and Σ′(*g*) to denote the set of automrphism of the representative GRN *g* after self-loop removal that only permutes Ω′, (b) *n*_*l*_(|Γ| − *k*) is the number of ways to allocate |Γ| − *k* labeled self-loops among the proteins Ω, (c) *m*_*s*_(*g*) is the number of ways to re-distribute the edges pointing from stimuli Ω_0_ in *g*, and (d) *r*(*g*) is keen to the number of ways to divide loci Γ into self-loops, non-self-loop edges pointing from stimuli, and others. As a result,

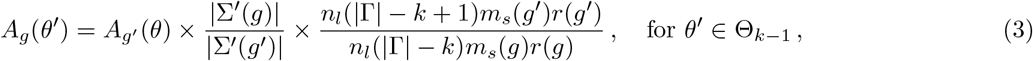

and it is noteworthy that *A*_*g*′_ (*θ*) is already known right after we constructed the layer Θ_*k*_ through Φ.

Third, we turn to the case where *θ*′ ∈ Θ_*k*_ but *θ*′ ≠ *θ*. Recall that, if any 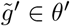 is a mutational neighbor of *g*, then there is a GRN 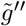 in layer Θ_*k*−1_ where 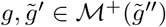, and such 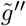 is unique up to arbitrary self-loop re-allocation. Additionally, the extra edge in *g* and 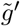 must corresponds to the same locus, so

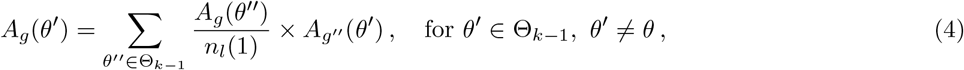

in which we abuse the analogy for *g*″ to be the representative GRN of equivalence class *θ*″. Lastly, if *θ*′ = *θ*, we also need to include the scenario that the mutational neighbor 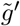 of *g* is generated by rewiring a self-loop to another self-loop. Therefore,

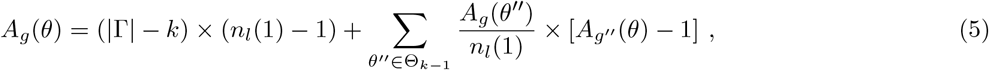

where the minus one in the square bracket avoids counting *g* itself multiple times.

In Algorithm 1, we summarize our proposed approach to construct the underlying space of a fitness landscape of GRNs through their equivalence classes Θ. It is apparent that the core of our algorithm is to find the joining operations Φ for a given layer Θ_*k*_. Fortunately, this task can be sufficiently achieved by pre-computing the phenotype-preserving automorphisms of every representative GRN once it is chosen. Suppose that we have revealed and stored the collection of joining operations for layers Θ_0_, Θ_1_, …, Θ_*k*−1_. Since these joining operations reflect the mutational neighbors and the phenotype-preserving isomorphisms in previous layers, the typed-(III) Φ for layer Θ_*k*_ are generated as compositions of the already uncovered operations. Furthermore, combining these stored joining operations and the newly computed automorphisms of representative GRNs in layer Θ_*k*_ gives rise to the remaining Φ of type (II). As a result, the only prerequisite in our proposed algorithm becomes to produce the phenotype-preserving automorphisms of a GRN.

## 4 Conclusion

In this work, we employ mechanistic knowledge of how phenotypes are computed from genotypes and regulatory interactions into fitness landscape models, where fitness is uniquely determined by the derived phenotype. This family of fitness landscape models features flexibility for tunable ruggedness and accessibility among phenotypes with bounded differences. Furthermore, we introduce the equivalence classes of GRNs, where GRNs of the same phenotype and with similar structural positions in the genotype network are coarse-grained into a group. These equivalence classes of GRNs lead to a compact and informative way to describe the fundamental space of a fitness landscape since the numerous GRNs in an equivalence class are simply mimicked by a single representative. We also develop an efficient approach to construct the underlying space of a fitness landscape based on the equivalences classes, where our method concurrently acquires the equivalence classes of GRNs and the mutational connection among the discovered classes.

### Algorithm 1

Constructing the underlying space of a fitness landscape of GRNs

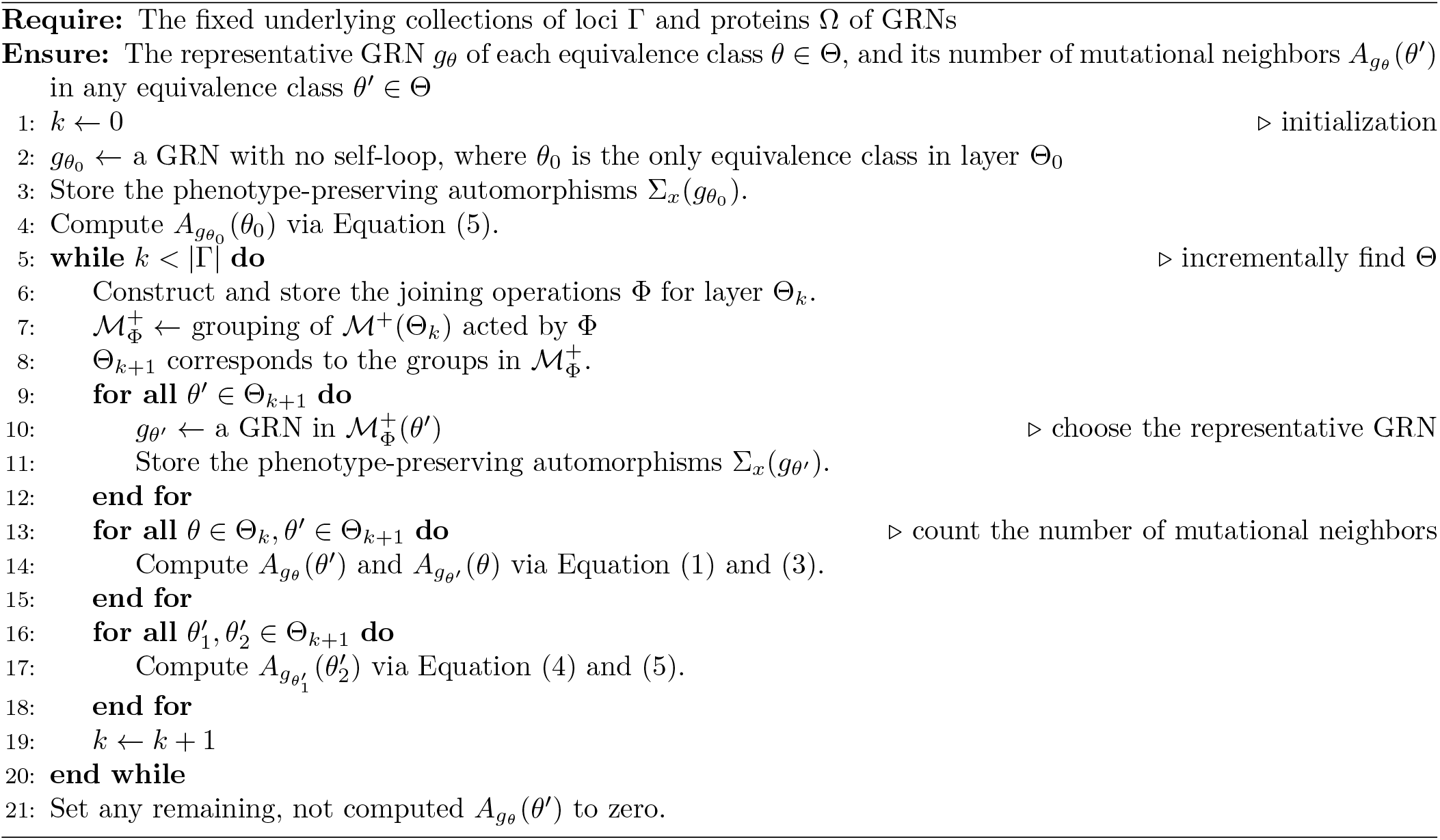

From a naive intuition, ruggedness and accessibility are two seemingly contradictory characteristics of a fitness landscape. Sign epistasis has been known to yield strong influence on a landscape’s ruggedness, and it was regarded as an impediment of evolutionary accessibility at its first introduction [18, 2]. On the contrary, recent studies proposed fitness landscape models that are keen to empirical observations and concluded that sign epistasis (and thus ruggedness) may co-exist with accessibility [54, 69]. Our model supports the latter for (a) the high dimensionality nature of GRNs [54], (b) that selection acts on the superposition of mutations and the background GRN rather than a few pairs of mutations [51], and perhaps more importantly, (c) that a GRN may experiences a series of neutral mutations and then evolve into a nearby phenotype [70, 7, 71, 3]. The discovered accessibility in a fitness landscape of GRNs hence highlights the role of neutral evolution the traditional metaphor, and it also agrees with the phenomenon of punctuated equilibrium [72, 36, 73].

Our derived equivalence classes of GRNs provide a novel, mesoscopic and optimally descriptive skeleton of a fitness landscape. As we mentioned in Introduction, neither of the genotypic space and the phenotypic space alone is best suited to characterize a fitness landscape. Advanced models incorporate the genotype-phenotype mapping and insert an addition phenotypic layer between genotypes and their fitness, where each individual genotype determines its phenotype (with a given environmental condition) and the phenotype specifies the fitness [59, 62, 63, 64]. These models are computationally expensive because their essence rests on exhausting all plausible genotypes to retain the necessary information. The equivalence classes of GRNs serves as an intermediate level between the genotypic and phenotypic space, and since they aggregate GRNs that share an common phenotype and alike mutational neighborhoods, the representative GRNs of these equivalence classes themselves are sufficient to compose the backbone of the fitness landscape.

We argue that the proposed constructive algorithm of a fitness landscape of GRNs is more efficient than the brute-force approach, where one enumerates all plausible GRNs and identifies the mutational neighbors for each of them. First of all, since we consolidate GRNs of an equivalence class into a single representative, our method is less costly in memory and it demands fewer computations when finding the mutational neighbors. Second, supposed all plausible GRNs are organized into layers by the number of non-self-loop edges (see section 3.3), it is noteworthy that every layer contains super-exponentially many GRNs. Our algorithm instead finds the equivalence classes in each layer iteratively: To construct the (*k* + 1)-th layer, we only have to exhaust the representative GRNs in the *k*-th layer along with any plausible, additional non-self-loop edge, whose amount is significantly fewer than the number of GRNs in the (*k* + 1)-th layer. In addition, existing heuristics for graph automorphisms [74, 75] can be extended to produce the phenotype-preserving automorphisms of the representative GRNs, which is the only prerequisite when joining together different GRN-edge pairs. Because the set of automorphisms becomes more limited as the complexity of a GRN increases, we expect a minor overhead in the joining procedure compared to the exhaustive, brute-force approach.

We envision the current work to aid future theoretical studies on evolutionary processes that take genotype-phenotype maps into account. The proposed mesoscopic backbone of a fitness landscape reduces complexity to enumerate the underlying genotypes, yet in a minimal way such that relations between genotypes and phenotypes are preserved so evolutionary processes modeled on the original landscape and the coarse-grained alternative are equivalent. The mesoscopic fitness landscape via equivalence classes of GRNs is hence more likely to generate theoretical predictions that was previously challenging. Furthermore, our methodology is applicable beyond GRNs to other models of genotype-phenotype mapping [76, 77], as long as the automorphisms of the genotype network are known. More broadly, this work showcases the potential to combine biological computation across different scales along the hierarchy, in particular computing biological functionality on the organism level and evolution on the population level.

## Acknowledgement

Hold.

## Funding

Hold.

## Conflicts of Interest

The authors declare no competing interests exist.

## Data Availability

The authors affirm that all data necessary for confirming the conclusions of the article are present within the article, figures, and tables.

## A Central and peripheral GRNs where no regulation presents

Here we show the connectivity of “central” and “peripheral” GRNs mentioned in section 3.1 under the extreme scenario where no gene regulation appears, in particular, when 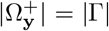. We observe that, since 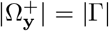, each node 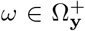 is incident to one and only one incoming edge, and thus each central GRN 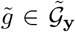 corresponds to a bijective mapping from 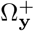 to edge labels Γ.

First, we show that if two central GRNs 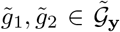 correspond to different mapping between 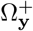 and Γ, then there is no mutational trajectory connecting them among 𝒢_**y**_. Start with assuming that such mutational trajectory does exist. Due to the different associated mappings of 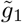 and 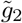, there is an 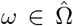 with distinct labels of the incident edge *γ*_1_ and *γ*_2_ in 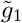 and 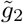 respectively. Because mutating 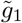 into 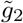 requires rewiring both *γ*_1_ and *γ*_2_, there must exist a GRN *g* where either none or both the edges *γ*_1_ and *γ*_2_ point to *ω*. Nevertheless, *g* contradicts with our observation following the constraint 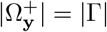. so *g* ∉ 𝒢_**y**_. As a result, under this extreme scenario, 𝒢_**y**_ fragment into multiple connected components when only mutations among themselves are considered.

Next, for any phenotype **y**′ for which 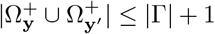, we show that there is a mutational trajectory among 𝒢_**y**_ connecting an arbitrary central GRN 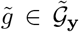 and a peripheral GRN *ĝ* ∈ 𝒢_**y**_ at the boundary of 𝒢_**y**_ and 𝒢_**y**′_. Specifically, take an 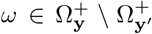 and its incident edge pointing from *ω*_0_ ∈ Ω_0_. For each 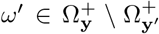 where *ω*′ ≠ *ω*, one can sequentially rewire the incident edge of *ω*′ to form a chain of 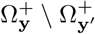 initiated by *ω*_0_, which lead to a resultant GRN *ĝ*. Moreover, since 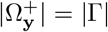, we have 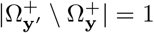. Recall from section 3.1, this *ĝ* is indeed peripheral GRN between 𝒢_**y**_ and 𝒢_**y**′_.

## B Phenotype-preserving automorphisms of the genotype network of GRNs

In this section we demonstrate (a) why GRNs mapped by automormphisms of the genotype network *G* are equivalent, (b) four graphical operations that generate phenotype-preserving automorphisms of *G*, and (c) the correctness of our iterative procedure to obtain a coarser parition than the equivalence classes of GRNs.

### Proposition B.1.

*Given an automorphism σ of the genotype network G and a mega-node function f*_*G*_ *that depends on the adjacency matrix* **A** *of G, for any GRNs g*_1_, *g*_2_ ∈ 𝒢 *where g*_2_ = *σ*(*g*_1_), *f*_*G*_(*g*_2_) = *f*_*G*_(*g*_1_).

*Proof*. Take 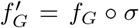. Since for any *g*_1_, *g*_2_ ∈ *V*(*G*), (*σ*(*g*_1_), *σ*(*g*_2_)) ∈ *E*(*G*) if and only if (*g*_1_, *g*_2_) ∈ *E*(*G*), the adjacency matrix *A* remains unchanged after permuting the mega-nodes through *σ*. As a result, we have 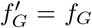, and 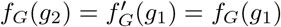.□

Let *π* and *π*′ be a permutation of the loci Γ and the dummy proteins Ω′ respectively. It is not hard to see that *π* and *π*′ also generate a permutation of the GRNs 𝒢. For *g* ∈ 𝒢, we abuse the notation *π*(*g*) to be the GRN mapped through the locus permutation *π*, where an edge with *e*_*g*_(*γ*) = (*u, v*) is transformed into that *e*_*π*(*g*)_(*π*(*γ*)) = (*u, v*). Similarly, in the GRN *π*′(*g*) mapped through the dummy protein permutation *π*′, an edge with *e*_*g*_(*γ*) = (*u, v*) is transformed into that *e*_*g*_(*γ*) = (*π*(*u*), *π*(*v*)).

Furthermore, we have two more types of graphical operations on GRNs:

### Definition B.1.

For a locus *γ* and two stimuli *ω, ω*′ ∈ Ω_0_, *ρ*_*γ,ω,ω*′_ : 𝒢 → 𝒢 transforms a GRN *g* into *g*′ such that edge *γ* becomes

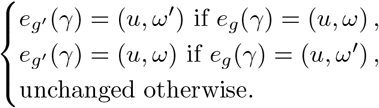

### Definition B.2.

For a locus *γ* and two node *ω* and *ω*′, *ϱ*_*γ,ω,ω*′_ : 𝒢 → 𝒢 transforms a GRN *g* into *g*′ such that edge *γ* becomes

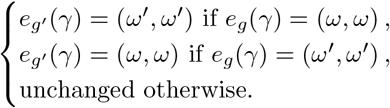

Note both *ρ*_*γ,ω,ω*′_ and *ϱ*_*γ,ω,ω*′_ are permutations of 𝒢 as well, where pairs of GRNs are mutually mapped from one to the other.

These four graphical operations introduced above are, more importantly, automorphisms of the genotype network *G* that also preserve the phenotype of GRNs:

### Theorem B.1.

*The transformations π, π*′, *ρ*_*γ,ω,ω*′_ *and ϱ*_*γ,ω,ω*′_ *are phenotype-preserving automorphisms of G*.

*Proof*. For *g*_1_, *g*_2_ ∈ 𝒢, let 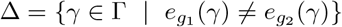. Since *π*′ is a permutation of dummy proteins, it preserves Δ. Additionally, because *ρ*_*γ,ω,ω*′_ and *ϱ*_*γ,ω,ω*′_ can be view as permutations of the source-target pair of a single edge *γ*, they also preserve Δ. The permutation *π* of Γ, on the other hand, does not preserve Δ, but it maintains its size |Δ|. Since (*g*_1_, *g*_2_) ∈ *E*(*G*) if and only if |Δ| = 1, the four transformations are automorphisms of *G*.

Furthermore, since *π, π*′ and *ϱ*_*γ,ω,ω*′_ simply changes the labels of edges, labels of the intermediate nodes, and the location of a self-loop, they maintain any path from a stimulus *ω*_0_ ∈ Ω_0_ to a fitness-relevant protein 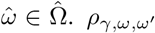 may alter the path between*ω*_0_ and 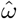 but the reachability from Ω_0_ to 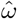 remains. Therefore, the four transformations also preserve the pheontype of GRNs.□

Finally, we turn to the computationally acquired partition that can be shown coarser than the equivalence classes of GRNs. Recall from section 3.2 that our iterative procedure starts from a partition 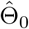 where GRNs with the same phenotype are grouped together. Given the partition 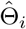, the next partition 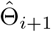 is obtained by further dividing groups in 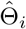 (if needed) such that for each group 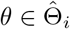 and 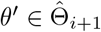, any two GRNs in *θ*′ have the same number of neighbors among *θ* in the genotype network *G*. The procedure terminates when a stationary partition 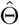 is reached. We then have:

### Theorem B.2.

*Every equivalence class θ* ∈ Θ *is included a group* 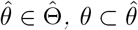.

*Proof*. Recall that GRNs in an equivalence class have the same phenotype, so for each *θ* ∈ Θ, there is some *θ*_0_ ∈ Θ_0_ where *θ* ⊂ *θ*_0_. Suppose that *θ* ⊂ *θ*_*i*_ for each *θ* ∈ Θ and some *θ*_*i*_ ∈ Θ_*i*_. Since Θ form an equitable partition, every *g* ∈ *θ* has the same number of neighbors in each *θ*′ ∈ Θ and thus also in each *θ*_*i*_ ∈ Θ_*i*_. Consequently, no two GRNs in *θ* will be separated into two different group in Θ_*i*+1_, and the theorem is proved by induction. □

## C Combining mutational neighbors into equivalence classes

In this section we tackle the two questions raised in section 3.3:

A. For an equivalence class *θ* ∈ Θ_*k*_ and its representative GRN *g* ∈ *θ*, under what condition will 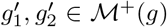 belong to the same equivalence class in layer Θ_*k*+1_?
B. For two distinct equivalence classes *θ*_1_, *θ*_2_ ∈ Θ_*k*_ and their representative GRNs *g*_1_ ∈ *θ*_1_ and *g*_2_ ∈ *θ*_2_, under what condition will 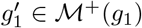 and 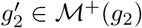 belong to the same equivalence class in layer Θ_*k*+1_?

And recall that for ease of demonstration we constrain the GRNs *g, g*_1_, *g*_2_, 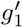 and 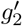 where only one stimulus node is incident to out-going edges.

### Definition C.1.

A phenotype-preserving isomorphism *π* from 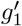 to 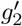 after self-loop removal is a permutation of loci Γ and dummy protein Ω′ such that for any locus *γ* and non-self-loop source-target pair 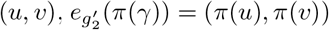 if and only if 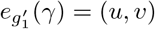.

Starting with question (A), we write 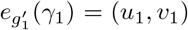 and 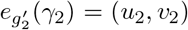 where *γ*_1_ and *γ*_2_ are the non-self-loop edge newly rewired to generate 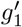 and 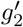 from *g* respectively. A few observations appear when we assume 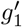 and 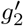 belong to the same equivalence class:

### Lemma C.1.

*Suppose a phenotype-preserving isomorphism π from* 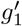 *to* 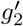 *after self-loop removal. There are two integers q* < *p such that π*^*p*^(*γ*_1_) = *γ*_1_, (*π*^*p*^(*u*_1_), *π*^*p*^(*v*_1_)) = (*u*_1_, *v*_1_), *and π*^*q*^(*γ*_1_) = *γ*_2_, (*π*^*q*^(*u*_1_), *π*^*q*^(*v*_1_)) = (*u*_2_, *v*_2_).

*Proof*. Since *π* is a permutation of finite sets, it must have a finite period, i.e., an integer *p* such that *π*^*p*^(*γ*_1_) = *γ*_1_ and (*π*^*p*^(*u*_1_), *π*^*p*^(*v*_1_)) = (*u*_1_, *v*_1_).

If *π*(*γ*_1_) = *γ*_2_ and (*π*(*u*_1_), *π*(*v*_1_)) = (*u*_2_, *v*_2_), then we have *q* = 1. Otherwise, it must map to a non-self-loop edge in *g* because *γ*_2_ is the only additional non-self-loop edge in *g*_2_, i.e., *e*_*g*_(*π*(*γ*_1_)) = (*π*(*u*_1_), *π*(*v*_1_)). Assume there is no integer *q* < *p* such that *π*^*q*^(*γ*_1_) = *γ*_2_ and (*π*^*q*^(*u*_1_), *π*^*q*^(*v*_1_)) = (*u*_2_, *v*_2_). Then *π*^*p*−1^(*γ*_1_) is a non-self-loop edge in *g* with *e*_*g*_(*π*^*p*−1^(*γ*_1_)) = (*π*^*p*−1^(*u*_1_), *π*^*p*−1^(*v*_1_)). However, since *γ*_1_ is not a non-self-loop edge in *g*_2_, the fact that *π*^*p*^(*γ*_1_) = *γ*_1_ and (*π*^*p*^(*u*_1_), *π*^*p*^(*v*_1_)) = (*u*_1_, *v*_1_) contradicts with *π* being an isomorphism from *g*_1_ to *g*_2_ after self-loop removal. Therefore, there is an integer *q* < *p* such that *π*^*q*^(*γ*_1_) = *γ*_2_ and (*π*^*q*^(*u*_1_), *π*^*q*^(*v*_1_)) = (*u*_2_, *v*_2_). □

### Lemma C.2.

*Suppose a phenotype-preserving isomorphism π from* 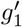 *to* 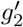 *after self-loop removal. For integers p and q in* *Lemma C*.*1*, 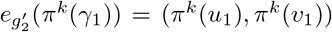 *for k* = 1, 2, …, *q, and* 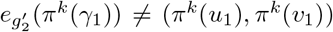 *for k* = *q* + 1, *q* + 2, …, *p*.

*Proof*. Since *π* is an isomorphism and *γ*_2_ = *π*^*q*^(*γ*_1_) is the only additional non-self-loop edge in *g*_2_, we have *π*(*γ*_1_), *π*^2^(*γ*_1_), …, *π*^*q*−1^(*γ*_1_) to be non-self-loop edges in *g*. Thus 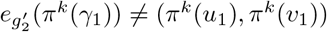 for *k* = 1, 2, …, *q*.On the other hand, since *γ*_2_ = *π*^*q*^(*γ*_1_) is not a non-self-loop edge in *g*_1_, the isomorphism *π* guaranteed that the source-target pairs (*π*^*q*+1^(*u*_1_), *π*^*q*+1^(*v*_1_)), (*π*^*q*+2^(*u*_1_), *π*^*q*+2^(*v*_1_)), …, (*π*^*p*^(*u*_1_), *π*^*p*^(*v*_1_)) do not match to edges in *g*_2_, in particular, 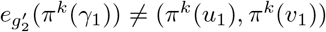 for *k* = *q* +1, *q* +2,…,*p*. □

### Lemma C.3.

*Suppose a phenotype-preserving isomorphism π from* 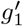 *to* 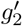 *after self-loop removal. Given integer q in* *Lemma C*.*1**, for a locus γ and non-self-loop source-target pair* (*u, v*) *where there is no* 0 ≤ *k* ≤ *q* − 1 *such that* (*γ, u, v*) = (*π*^*k*^(*γ*), *π*^*k*^(*u*_1_), *π*^*k*^(*v*_1_)), *e*_*g*_(*π*(*γ*)) = (*π*(*u*), *π*(*v*)) *if and only if e*_*g*_(*γ*) = (*u, v*).

*Proof*. For 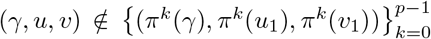, since (*γ*_1_, *u*_1_, *v*_1_) and (*γ*_2_, *u*_2_, *v*_2_) are already excluded and by Definition C.1, we have *e*_*g*_(*π*(*γ*)) = (*π*(*u*), *π*(*v*)) if and only if *e*_*g*_(*γ*) = (*u, v*). Furthermore, for *k* = *q, q* + 1, …, *p* − 1, because the only additional non-self-loop edge in 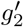 follows *π*^*q*^(*γ*_1_) = *γ*_2_ and (*π*^*q*^(*u*_1_), *π*^*q*^(*v*_1_)) = (*u*_2_, *v*_2_), according to Lemma C.2, we know that *e*_*g*_(*π*^*k*^(*γ*_1_)) ≠ (*π*^*k*^(*u*_1_), *π*^*k*^(*v*_1_)) and *e*_*g*_(*π*^*k*+1^(*γ*_1_)) ≠ (*π*^*k*+1^(*u*_1_), *π*^*k*+1^(*v*_1_)). As a result, the statement in Lemma C.3 is true for any locus *γ* and non-self-loop source-target pair (*u, v*) where 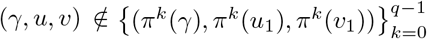.□

The following theorem resolves the necessary and sufficient condition in question (A):

### Theorem C.1.

*Let* 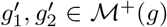 *with the additional non-self-loop edge* 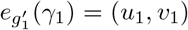 *and* 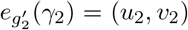 *respectively*. 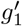 *and* 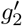 *belong to the same equivalence class if and only if there exist two integers q* < *p and an phenotype-preserving automorphism σ of a subgraph* 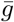 *of g such that*

i. (*σ*^*q*^(*γ*_1_), *σ*^*q*^(*u*_1_), *σ*^*q*^) = (*γ*_2_, *u*_2_, *v*_2_),
ii. (*σ*^*p*^(*γ*_1_), *σ*^*p*^(*u*_1_), *σ*^*p*^(*v*_1_)) = (*γ*_1_, *u*_1_, *v*_1_),
iii. 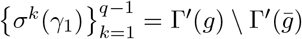 *and e*_*g*_(*σ*^*k*^(*γ*_*1*_)) = (*σ*^*k*^(*u*_1_), *σ*^*k*^(*v*_1_) *for k* = 1, 2, …, *q* − 1, *and*
iv. *e*_*g*_ *σ*^*k*^(*γ*_1_) ≠ *σ*^*k*^(*u*_1_), *σ*^*k*^(*v*_1_) *for k* = *q, q* + 1, …, *p*.

*Proof*. For one direction, suppose a phenotype-preserving isomorphism *π* from 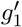 to 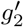 after self-loop removal. According to Lemma C.1–C.3, we see that *π* is a phenotype-preserving automorphism of a subgraph 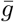 of *g* where 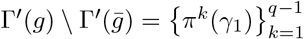. Taking *σ* = *π* satisfies all the four conditions.

For the other direction, we show that such a phenotype-preserving automorphism *σ* is also a phenotype-preserving isomorphism from 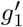 to 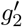 after self-loop removal. First, regarding any locus *γ* and non-self-loop target pair (*u, v*) such that 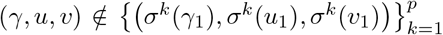, we have 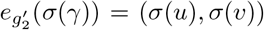 if and only if 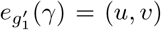 because *σ* is an automorphism of 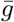 and *γ*_1_, *γ*_2_ and 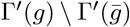 are already excluded. Second, for *k* = 0, 1, …, *q* − 1, the conditions i.–iii. ensure that 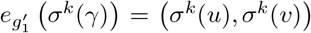 and 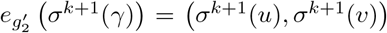. Lastly, for *k* = *q, q* + 1, …, *p* − 1, the conditions i., ii. and iv. indicates 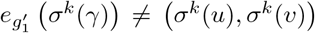 and 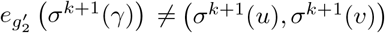. □

Next, the necessary and sufficient condition of question (B) is described in the theorem below:

### Theorem C.2.

*Let g*_1_ *and g*_2_ *be of two different equivalence classes, and let* 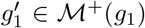*and* 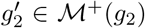 *with the additional non-self-loop edge* 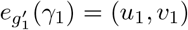 *and* 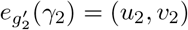 *respectively*. 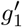 *and* 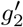 *belong to the same equivalence class if and only if there exist GRNs* 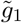 *and g*″ *such that*

i. 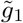*and g*_1_ *belong to the same equivalence class*,
ii. 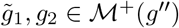, *and*
iii. 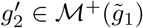.

*Proof*. For one direction, suppose a phenotype-preserving isomorphism *π* from 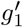 to 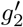 after self-loop removal. Take 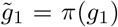, i.e., 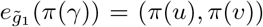 for each *γ* ∈ Γ′(*g*_1_) with 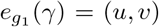, so *π* also becomes a phenotype-preserving isomorphism from *g*_1_ to 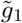. Moreover, due to the isomorphism *π*, we note the 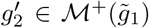 with the additional non-self-loop edge 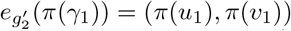. Let *g*″ be a GRN obtained by rewiring edges *π*(*γ*_1_) and *γ*_2_ from 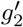 to two arbitrary self-loops. We have 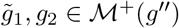. with the additional non-self-loop edge 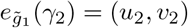 and 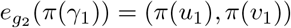 respectively. Observe that the GRNs 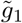 and *g*″ satisfy the conditions i.–iii..

For the other direction, suppose there exist GRNs 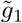 and *g*″ where the conditions i.–iii. hold. Let *π* be the phenotype-preserving isomorphism from *g*_1_ to 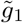 after self-loop removal such that the additional non-self-loop edge of *g*_2_ ∈ ℳ^+^(*g*″) complies 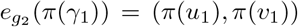. Since 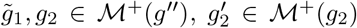 and 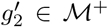, we also have the addition non-self-loop edge in 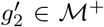 to follow 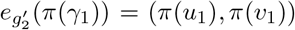. Therefore, *π* is also a phenotype-preserving isomorphism from 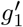 to 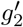 after self-loop removal. □

## D Size of an equivalence class of GRNs

Here, given the representative GRN *g* of an equivalence class *θ*, we calculate the the number of GRNs in *θ* in Equation 2. Observe that |*θ*| is proportional to (a) the number of GRNs to which there is a phenotype-preserving isomorphism from *g*, (b) the number of ways to arbitrarily allocate self-loops on *g*, and (c) the number of ways to arbitrarily rewire source nodes of edges among the stimuli Ω_0_.

First, for part (a), if we temporally ignore labels on the edges, then the set of permutations over dummy proteins Ω′ is partitioned into groups of isomorphisms from *g* to different GRNs. In addition, the size of each group is exactly the number of automorphisms of *g*, since compositing an automorphism of *g* and an isomorphism from *g* to *g*′ also generates an isomorphism from *g* to *g*′. Thus there are in total 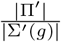 different GRNs isomorphic to *g*, where Π′ is the set of permutations over Ω′ and Σ′(*g*) is the set of automorphisms of *g* that only permutes Ω′.

Second, for part (b), every possible allocation distributes |Γ| − |Γ′(*g*)| self-loops over |Ω \ Ω_0_| nodes. Supposed that the labels of self-loops Γ \ Γ′(*g*) are given, we have

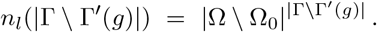

Third, we write *k*_*s*_(*g*) as the number of incident edges to stimuli Ω_0_ in *g*. Any possibility in part (c) chooses a source node among Ω_0_ for each of the *k*_*s*_(*g*) incident edges. Providing that the labels of incident edges of the stimuli are already known, we have

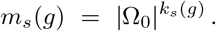

Combined, the size of the equivalence class *θ* becomes

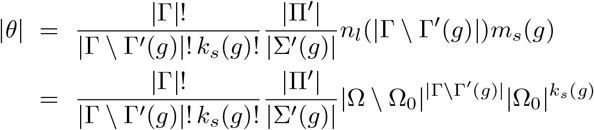

where the first fraction represents first selecting combinations of |Γ\Γ′(*g*) | and *k*_*s*_(*g*) labels for self-loops and edges incident to stimuli, and then permuting the remaining labels, which also contributes to different phenotype-preserving isomorphisms from *g* but was previously omitted.

## Footnotes

1 In this paper, we assume that the stimuli Ω_0_ must be proteins which can not be produced by expression, and we leave no constraint to the fitness-relevant and dummy proteins 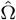 and Ω ′.

2 Particularly, this is an edge outside of a spanning tree of central GRN 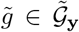.

3 A permutation of 𝒢 is a mapping *σ* : 𝒢 → 𝒢 where no two GRNs are mapped to the same GRN, i.e., *σ*(*g*_1_) ≠ *σ*(*g*_2_) if *g*_1_ ≠ *g*_2_ for any *g*_1_, *g*_2_ ∈ 𝒢.

4 A partition 𝒫 of elements *X* is a set of sets/groups 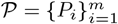 where each *x* ∈ *X* belongs to one and only one group.

5 A partition 𝒫 is coarser than another partition 𝒫′ if any group in 𝒫′ is included in some group in 𝒫.

6 Constraining on that the isomorphism *π* preserves phenotype, we note *π* must map each fitness-relevant protein to itself.

## Notes

### Competing Interest Statement

The authors have declared no competing interest.

## References

[1] Sewall Wright. The roles of mutation, inbreeding, crossbreeding, and selection in evolution. Proceedings of the Sixth International Congress on Genetics, pages 356–366, 1932.

[2] J Arjan Gm De Visser and Joachim Krug. Empirical fitness landscapes and the predictability of evolution. Nature Reviews Genetics, 15(7):480–490, 2014.

[3] Inês Fragata, Alexandre Blanckaert, Marco António Dias Louro, David A Liberles, and Claudia Bank. Evolution in the light of fitness landscape theory. Trends in ecology & evolution, 34(1):69–82, 2019.

[4] Ivan G Szendro, Martijn F Schenk, Jasper Franke, Joachim Krug, and J Arjan GM De Visser. Quantitative analyses of empirical fitness landscapes. Journal of Statistical Mechanics: Theory and Experiment, 2013(01):P01005, 2013.

[5] Kavita Jain and Joachim Krug. Deterministic and stochastic regimes of asexual evolution on rugged fitness landscapes. Genetics, 175(3):1275–1288, 2007.

[6] Sergey Kryazhimskiy, Gašper Tkačik, and Joshua B Plotkin. The dynamics of adaptation on correlated fitness landscapes. Proceedings of the National Academy of Sciences, 106(44):18638–18643, 2009.

[7] Jeremy A Draghi, Todd L Parsons, Günter P Wagner, and Joshua B Plotkin. Mutational robustness can facilitate adaptation. Nature, 463(7279):353–355, 2010.

[8] Nicholas C Wu, Lei Dai, C Anders Olson, James O Lloyd-Smith, and Ren Sun. Adaptation in protein fitness landscapes is facilitated by indirect paths. Elife, 5:e16965, 2016.

[9] Sergey Gavrilets. Fitness landscapes and the origin of species. Princeton University Press, 2004.

[10] Christelle Fraïsse, P Alexander Gunnarsson, Denis Roze, Nicolas Bierne, and John J Welch. The genetics of speciation: insights from fisher’s geometric model. Evolution, 70(7):1450–1464, 2016.

[11] J Arjan GM de Visser, Su-Chan Park, and Joachim Krug. Exploring the effect of sex on empirical fitness landscapes. the american naturalist, 174(S1):S15–S30, 2009.

[12] Sarah P Otto. The evolutionary enigma of sex. the american naturalist, 174(S1):S1–S14, 2009.

[13] Richard A Watson, Daniel M Weinreich, and John Wakeley. Genome structure and the benefit of sex. Evolution: International Journal of Organic Evolution, 65(2):523–536, 2011.

[14] H Allen Orr. The genetic theory of adaptation: a brief history. Nature Reviews Genetics, 6(2):119–127, 2005.

[15] Alexander E Lobkovsky, Yuri I Wolf, and Eugene V Koonin. Predictability of evolutionary trajectories in fitness landscapes. PLoS Comput Biol, 7(12):e1002302, 2011.

[16] Merijn LM Salverda, Eynat Dellus, Florien A Gorter, Alfons JM Debets, John Van Der Oost, Rolf F Hoekstra, Dan S Tawfik, and J Arjan GM de Visser. Initial mutations direct alternative pathways of protein evolution. PLoS Genet, 7(3):e1001321, 2011.

[17] Claudia Bank, Sebastian Matuszewski, Ryan T Hietpas, and Jeffrey D Jensen. On the (un) predictability of a large intragenic fitness landscape. Proceedings of the National Academy of Sciences, 113(49):14085–14090, 2016.

[18] Daniel M Weinreich, Nigel F Delaney, Mark A DePristo, and Daniel L Hartl. Darwinian evolution can follow only very few mutational paths to fitter proteins. science, 312(5770):111–114, 2006.

[19] Hsin-Hung Chou, Hsuan-Chao Chiu, Nigel F Delaney, Daniel Segré, and Christopher J Marx. Diminishing returns epistasis among beneficial mutations decelerates adaptation. Science, 332(6034):1190–1192, 2011.

[20] Aisha I Khan, Duy M Dinh, Dominique Schneider, Richard E Lenski, and Tim F Cooper. Negative epistasis between beneficial mutations in an evolving bacterial population. Science, 332(6034):1193–1196, 2011.

[21] JAGM De Visser, Rolf F Hoekstra, and Herman van den Ende. Test of interaction between genetic markers that affect fitness in aspergillus niger. Evolution, 51(5):1499–1505, 1997.

[22] David W Hall, Matthew Agan, and Sara C Pope. Fitness epistasis among 6 biosynthetic loci in the budding yeast saccharomyces cerevisiae. Journal of Heredity, 101(Suppl 1):S75–S84, 2010.

[23] Michael C Whitlock and Denis Bourguet. Factors affecting the genetic load in drosophila: synergistic epistasis and correlations among fitness components. Evolution, 54(5):1654–1660, 2000.

[24] Trevor Hinkley, Jõao Martins, Colombe Chappey, Mojgan Haddad, Eric Stawiski, Jeannette M Whitcomb, Christos J Petropoulos, and Sebastian Bonhoeffer. A systems analysis of mutational effects in hiv-1 protease and reverse transcriptase. Nature genetics, 43(5):487–489, 2011.

[25] Roger D Kouyos, Gabriel E Leventhal, Trevor Hinkley, Mojgan Haddad, Jeannette M Whitcomb, Christos J Petropoulos, and Sebastian Bonhoeffer. Exploring the complexity of the hiv-1 fitness landscape. PLoS Genet, 8(3):e1002551, 2012.

[26] Ryan T Hietpas, Jeffrey D Jensen, and Daniel NA Bolon. Experimental illumination of a fitness landscape. Proceedings of the National Academy of Sciences, 108(19):7896–7901, 2011.

[27] Jakub Otwinowski and Ilya Nemenman. Genotype to phenotype mapping and the fitness landscape of the e. coli lac promoter. PloS one, 8(5):e61570, 2013.

[28] Karen S Sarkisyan, Dmitry A Bolotin, Margarita V Meer, Dinara R Usmanova, Alexander S Mishin, George V Sharonov, Dmitry N Ivankov, Nina G Bozhanova, Mikhail S Baranov, Onuralp Soylemez, et al. Local fitness landscape of the green fluorescent protein. Nature, 533(7603):397–401, 2016.

[29] Zoë N Rogers, Christopher D McFarland, Ian P Winters, Jose A Seoane, Jennifer J Brady, Stephanie Yoon, Christina Curtis, Dmitri A Petrov, and Monte M Winslow. Mapping the in vivo fitness landscape of lung adenocarcinoma tumor suppression in mice. Nature genetics, 50(4):483–486, 2018.

[30] Caroline J Watson, AL Papula, Gladys YP Poon, Wing H Wong, Andrew L Young, Todd E Druley, Daniel S Fisher, and Jamie R Blundell. The evolutionary dynamics and fitness landscape of clonal hematopoiesis. Science, 367(6485):1449–1454, 2020.

[31] William Rowe, Mark Platt, David C Wedge, Philip J Day, Douglas B Kell, and Joshua Knowles. Analysis of a complete dna–protein affinity landscape. Journal of The Royal Society Interface, 7(44):397–408, 2010.

[32] Jason N Pitt and Adrian R Ferré-D’Amaré. Rapid construction of empirical rna fitness landscapes. Science, 330(6002):376–379, 2010.

[33] José I Jiménez, Ramon Xulvi-Brunet, Gregory W Campbell, Rebecca Turk-MacLeod, and Irene A Chen. Comprehensive experimental fitness landscape and evolutionary network for small rna. Proceedings of the National Academy of Sciences, 110(37):14984–14989, 2013.

[34] Chuan Li, Wenfeng Qian, Calum J Maclean, and Jianzhi Zhang. The fitness landscape of a trna gene. Science, 352(6287):837–840, 2016.

[35] Devin P Bendixsen, Bjørn østman, and Eric J Hayden. Negative epistasis in experimental rna fitness landscapes. Journal of molecular evolution, 85(5):159–168, 2017.

[36] Jacobo Aguirre, Pablo Catalán, José A Cuesta, and Susanna Manrubia. On the networked architecture of genotype spaces and its critical effects on molecular evolution. Open Biology, 8(7):180069, 2018.

[37] John FC Kingman. A simple model for the balance between selection and mutation. Journal of Applied Probability, 15(1):1–12, 1978.

[38] Stuart Kauffman and Simon Levin. Towards a general theory of adaptive walks on rugged landscapes. Journal of theoretical Biology, 128(1):11–45, 1987.

[39] Stuart A Kauffman and Edward D Weinberger. The nk model of rugged fitness landscapes and its application to maturation of the immune response. Journal of theoretical biology, 141(2):211–245, 1989.

[40] Takuyo Aita, Hidefumi Uchiyama, Tetsuya Inaoka, Motowo Nakajima, Toshio Kokubo, and Yuzuru Husimi. Analysis of a local fitness landscape with a model of the rough mt. fuji-type landscape: Application to prolyl endopeptidase and thermolysin. Biopolymers: Original Research on Biomolecules, 54(1):64–79, 2000.

[41] Johannes Neidhart, Ivan G Szendro, and Joachim Krug. Adaptation in tunably rugged fitness landscapes: the rough mount fuji model. Genetics, 198(2):699–721, 2014.

[42] Olivier Tenaillon. The utility of fisher’s geometric model in evolutionary genetics. Annual review of ecology, evolution, and systematics, 45:179–201, 2014.

[43] Ronald A Fisher. The general theory of natural selection. Oxford University Press, 1930.

[44] Takuyo Aita, Masahiro Iwakura, and Yuzuru Husimi. A cross-section of the fitness landscape of dihydrofolate reductase. Protein Engineering, 14(9):633–638, 2001.

[45] Maurício Carneiro and Daniel L Hartl. Adaptive landscapes and protein evolution. Proceedings of the National Academy of Sciences, 107(Suppl 1):1747–1751, 2010.

[46] Frank J Poelwijk, Sorin Tănase-Nicola, Daniel J Kiviet, and Sander J Tans. Reciprocal sign epistasis is a necessary condition for multi-peaked fitness landscapes. Journal of Theoretical Biology, 272(1):141–144, 2011.

[47] Elena R Lozovsky, Thanat Chookajorn, Kyle M Brown, Mallika Imwong, Philip J Shaw, Sumalee Kamchon-wongpaisan, Daniel E Neafsey, Daniel M Weinreich, and Daniel L Hartl. Stepwise acquisition of pyrimethamine resistance in the malaria parasite. Proceedings of the National Academy of Sciences, 106(29):12025–12030, 2009.

[48] Mark Lunzer, Stephen P Miller, Roderick Felsheim, and Antony M Dean. The biochemical architecture of an ancient adaptive landscape. Science, 310(5747):499–501, 2005.

[49] Jamie T Bridgham, Sean M Carroll, and Joseph W Thornton. Evolution of hormone-receptor complexity by molecular exploitation. Science, 312(5770):97–101, 2006.

[50] Frank J Poelwijk, Daniel J Kiviet, and Sander J Tans. Evolutionary potential of a duplicated repressor-operator pair: simulating pathways using mutation data. PLoS Computational Biology, 2(5):e58, 2006.

[51] Frank J Poelwijk, Daniel J Kiviet, Daniel M Weinreich, and Sander J Tans. Empirical fitness landscapes reveal accessible evolutionary paths. Nature, 445(7126):383–386, 2007.

[52] Jasper Franke, Alexander Klözer, J Arjan GM de Visser, and Joachim Krug. Evolutionary accessibility of mutational pathways. PLoS Compututational Biology, 7(8):e1002134, 2011.

[53] Peter Hegarty, Anders Martinsson, et al. On the existence of accessible paths in various models of fitness landscapes. Annals of Applied Probability, 24(4):1375–1395, 2014.

[54] Marcin Zagorski, Zdzislaw Burda, and Bartlomiej Waclaw. Beyond the hypercube: evolutionary accessibility of fitness landscapes with realistic mutational networks. PLoS Computational Biology, 12(12):e1005218, 2016.

[55] Douglas B Kell. Genotype–phenotype mapping: genes as computer programs. TRENDS in Genetics, 18(11): 555–559, 2002.

[56] Hue Sun Chan and Erich Bornberg-Bauer. Perspectives on protein evolution from simple. Applied Bioinformatics, 50(3):121–144, 2002.

[57] Michael Stich, Ester Lázaro, and Susanna C Manrubia. Phenotypic effect of mutations in evolving populations of rna molecules. BMC Evolutionary Biology, 10(1):1–17, 2010.

[58] Michael E Palmer, Arnav Moudgil, and Marcus W Feldman. Long-term evolution is surprisingly predictable in lattice proteins. Journal of The Royal Society Interface, 10(82):20130026, 2013.

[59] Shimon Bershtein, Adrian WR Serohijos, and Eugene I Shakhnovich. Bridging the physical scales in evolutionary biology: from protein sequence space to fitness of organisms and populations. Current opinion in structural biology, 42:31–40, 2017.

[60] Lilia Perfeito, Stephane Ghozzi, Johannes Berg, Karin Schnetz, and Michael Lässig. Nonlinear fitness landscape of a molecular pathway. PLoS Genetics, 7(7):e1002160, 2011.

[61] Hsin-Hung Chou, Nigel F Delaney, Jeremy A Draghi, and Christopher J Marx. Mapping the fitness landscape of gene expression uncovers the cause of antagonism and sign epistasis between adaptive mutations. PLoS Genetics, 10(2):e1004149, 2014.

[62] Tamar Friedlander, Roshan Prizak, Nicholas H Barton, and Gašper Tkačik. Evolution of new regulatory functions on biophysically realistic fitness landscapes. Nature Communications, 8(1):1–11, 2017.

[63] Thomas D Cuypers, Jacob P Rutten, and Paulien Hogeweg. Evolution of evolvability and phenotypic plasticity in virtual cells. BMC Evolutionary Biology, 17(1):1–16, 2017.

[64] Pablo Yubero, Susanna Manrubia, and Jacobo Aguirre. The space of genotypes is a network of networks: implications for evolutionary and extinction dynamics. Scientific Reports, 7(1):1–12, 2017.

[65] Noémie Harmand, Romain Gallet, Roula Jabbour-Zahab, Guillaume Martin, and Thomas Lenormand. Fisher’s geometrical model and the mutational patterns of antibiotic resistance across dose gradients. Evolution, 71(1):23–37, 2017.

[66] Chia-Hung Yang and Samuel V Scarpino. Reproductive barriers as a byproduct of gene network evolution. bioRxiv, 2020.

[67] Chia-Hung Yang and Samuel V Scarpino. The ensemble of gene regulatory networks at mutation-selection balance. bioRxiv, 2021.

[68] Chris D Godsil. Compact graphs and equitable partitions. Linear Algebra and its Applications, 255(1-3):259–266, 1997.

[69] Suman G Das, Susana OL Direito, Bartlomiej Waclaw, Rosalind J Allen, and Joachim Krug. Predictable properties of fitness landscapes induced by adaptational tradeoffs. Elife, 9:e55155, 2020.

[70] Andreas Wagner. Neutralism and selectionism: a network-based reconciliation. Nature Reviews Genetics, 9(12):965–974, 2008.

[71] Devin P Bendixsen, James Collet, Bjørn østman, and Eric J Hayden. Genotype network intersections promote evolutionary innovation. PLoS Biology, 17(5):e3000300, 2019.

[72] Gene Hunt, Melanie J Hopkins, and Scott Lidgard. Simple versus complex models of trait evolution and stasis as a response to environmental change. Proceedings of the National Academy of Sciences, 112(16):4885–4890, 2015.

[73] Lydia R Heasley, Nadia MV Sampaio, and Juan Lucas Argueso. Systemic and rapid restructuring of the genome: a new perspective on punctuated equilibrium. Current Genetics, pages 1–7, 2020.

[74] José Luis López-Presa, Luis F Chiroque, and Antonio Fernández Anta. Novel techniques to speed up the computation of the automorphism group of a graph. Journal of Applied Mathematics, 2014, 2014.

[75] Stoicho D Stoichev. New exact and heuristic algorithms for graph automorphism group and graph isomorphism. Journal of Experimental Algorithmics (JEA), 24:1–27, 2019.

[76] Ting Hu, Marco Tomassini, and Wolfgang Banzhaf. A network perspective on genotype–phenotype mapping in genetic programming. Genetic Programming and Evolvable Machines, 21(3):375–397, 2020.

[77] Sam F Greenbury, Ard A Louis, and Sebastian E Ahnert. The structure of genotype-phenotype maps makes fitness landscapes navigable. bioRxiv, 2021.

